# Sequence memory in recurrent neuronal network can develop without structured input

**DOI:** 10.1101/2020.09.15.297580

**Authors:** Matthias Loidolt, Lucas Rudelt, Viola Priesemann

## Abstract

How does spontaneous activity during development prepare cortico-cortical connections for sensory input? We here analyse the development of sequence memory, an intrinsic feature of recurrent networks that supports temporal perception. We use a recurrent neural network model with homeostatic and spike-timing-dependent plasticity (STDP). This model has been shown to learn specific sequences from structured input. We show that development even under unstructured input increases unspecific sequence memory. Moreover, networks “pre-shaped” by such unstructured input subsequently learn specific sequences faster. The key structural substrate is the emergence of strong and directed synapses due to STDP and synaptic competition. These construct self-amplifying preferential paths of activity, which can quickly encode new input sequences. Our results suggest that memory traces are not printed on a *tabula rasa*, but instead harness building blocks already present in the brain.

## Introduction

Distributed computation, as performed by neurons in the mammalian brain, needs some form of memory. In particular, sequence memory is necessary for control of complex movement and perception of temporal stimuli. Sequence memory has been observed in cortical areas as diverse as inferior temporal cortex [Meyer et al., 2014], extrastriate cortex [Eagleman and Dragoi, 2012] or primary visual cortex [Gavornik and Bear, 2014]. The capabilities for sequence memory arise early in human development [Mandel et al., 1996]. We here ask if the capabilities for sequence memory can develop even before the onset of sensory stimuli, specifically through the interaction of neural plasticity and spontaneous activity.

Sequence memory is the ability to recognize an input sequence from its first elements [Gavornik and Bear, 2014]. Biophysically, this sequence memory can be implemented by the “echo state” or”fading memory” property [Maass et al., 2002] which is inherent to the dynamics of recurrently connected neural networks [Nikolić et al., 2009, Ju et al., 2015]: the recurrent connections project temporal dynamics of the input into the high-dimensional dynamics of the network activity. This high dimensionality allows past input to influence future activity, and this past input can be inferred from current activity. Thereby the network activity provides a form of “active” or “dynamic” sequence memory [Boedecker et al., 2012]. For specific sequences, this “dynamic” memory can be optimized through the process of sequence learning [Lazar et al., 2009, Wang et al., 2017]. Sequence learning requires multiple exposures to the same sequence in order to adapt recurrent connections to input properties via neural plasticity.

Neural plasticity mechanisms shape networks during development [Turrigiano and Nelson, 2004] and enable learning in adulthood [Biane et al., 2019]: activity-dependent plasticity rules, such as Hebbian learning [Hebb, 1949] and spike-timing-dependent-plasticity (STDP) [Bi and Poo, 1998], establish useful connections based on the correlations of pre- and postsynaptic spiking activity. Homeostatic plasticity rules, such as Synaptic Scaling/Synaptic Normalisation (SN) [Wallace and Bear, 2004] and Intrinsic Plasticity (IP) [Echegoyen et al., 2007] adapt neuronal excitability and re-distribute synaptic efficacies to regulate the overall spiking activity. *In vivo*, these plasticity rules presumably enable surprisingly fast sequence learning: only tens of presentations suffice to establish memory [Brea and Gerstner, 2016, Kang et al., 2017]. *In silico* models implementing realistic timescales and strengths of neural plasticity, on the contrary, require thousands of presentations to establish memory [Brea and Gerstner, 2016].

This discrepancy could be resolved by making use of spontaneous activity, which is ubiquitous in the nervous system during development and adulthood [Chiu et al., 2004]. Even before eye opening, spontaneous activity is generated in the sensory periphery and cortex proper [Luhmann et al., 2016]. There is exhaustive evidence that periphery-generated, structured spontaneous activity refines feed-forward connections [Luhmann et al., 2016, Richter and Gjorgjieva, 2017]. Yet, the effects of cortically-generated, unstructured spontaneous activity on cortico-cortical connections remain largely unknown [Richter and Gjorgjieva, 2017].

We here ask if neural networks subject to simple, unstructured spontaneous activity can develop sequence memory even before the onset of structured sensory input. To this end, we use an established recurrent network model of sequence learning (SORN, Lazar et al. [2009], Wang et al. [2017]) that is equipped with spike-timing dependent plasticity (STDP) and homeostatic plasticity rules on two different timescales, and has been shown to reproduce central features of cortical spiking dynamics [Hartmann et al., 2015]. We then show that development in the presence of spontaneous activity, modeled as unstructured noise [Triplett et al., 2018, Zierenberg et al., 2018], improves sequence memory in an unspecific manner. Furthermore, we show that this improvement is maintained even when the network continues development under sequence input, speeding up consecutive learning of specific sequences. Thus, spontaneous activity during development can pre-shape the network towards strongly improved performance when the animal encounters the first structured sequences. We provide a mechanistic understanding of the improved sequence memory: preferential paths of few strong and directed synapses arise from initial variances in the synaptic matrix that are amplified by STDP and synaptic competition. Finally, we conclude that these can be harnessed to quickly form new memory traces.

## Methods Summary

### Network Model

To ensure comparability, we use exactly the same network model and parameters as the SORN introduced in Lazar et al. [2009]. In brief, we use a recurrent network with *N*^*E*^ = 200 excitatory and *N*^*I*^ = 0.2 × *N*^*E*^ = 40 inhibitory binary threshold neurons (Fig. 1A). The sequence memory mechanism we propose shows qualitatively the same development in larger networks (*N*^*E*^ = 1000; Fig. 3-S4). At every discrete timestep *t* (of size Δ*t* = 10 ms), the activity of excitatory neurons 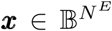 and inhibitory neurons 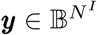 is given by:

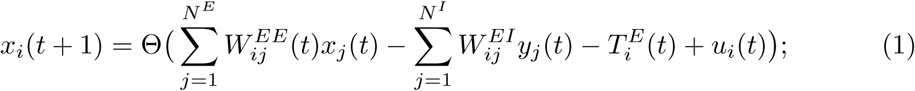

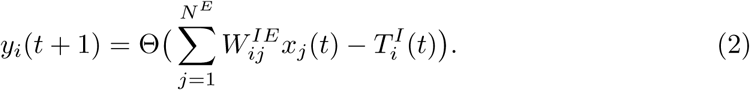

**Figure 1.**
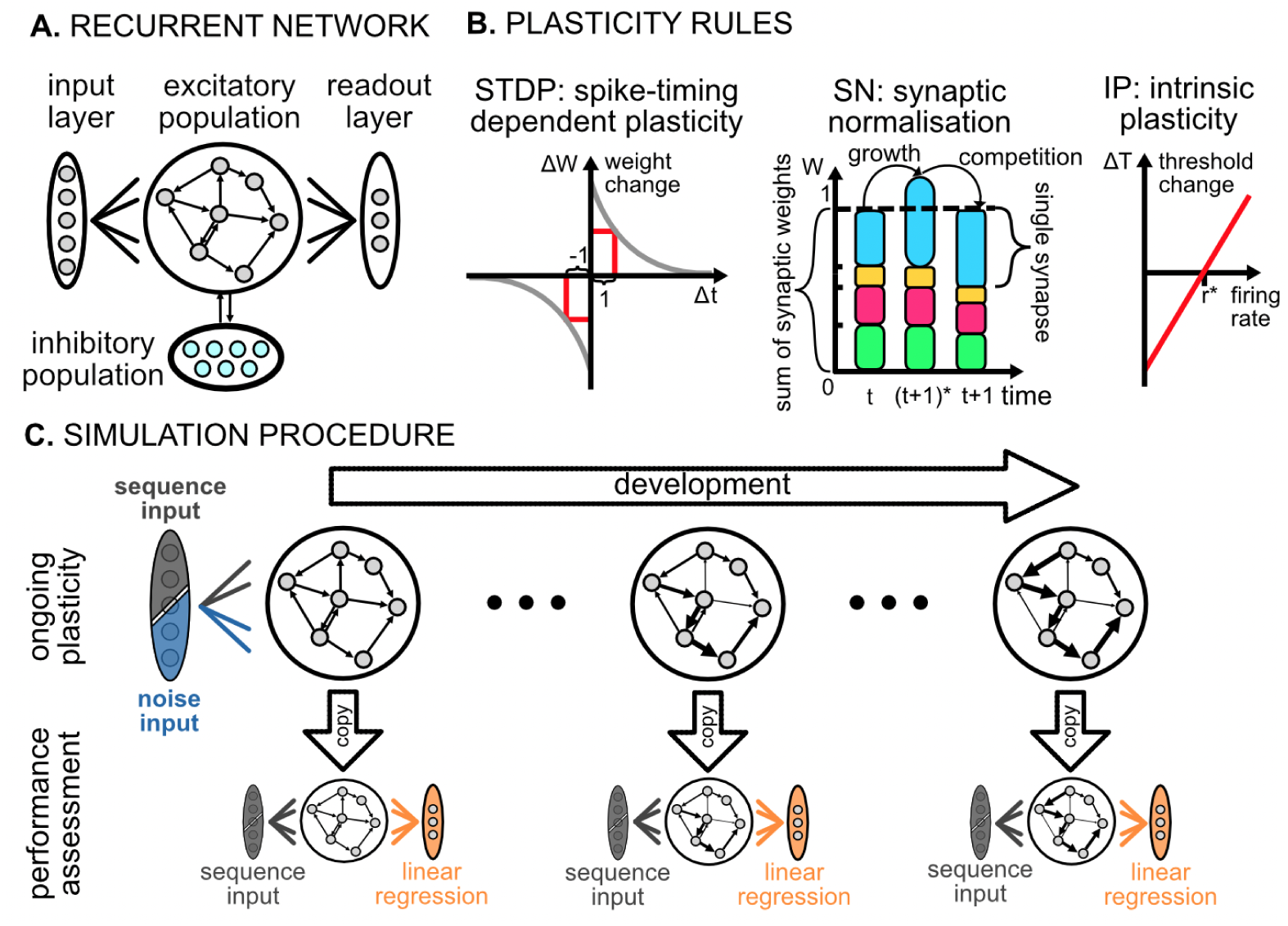
Recurrent network model and simulation of development under different input conditions. **A**. Sketch of the recurrent network model with excitatory and inhibitory populations of binary threshold neurons that are fully connected to each other. Excitatory-excitatory connections are sparse, recurrent, and subject to plasticity. Inhibitory-inhibitory connections are nil. The excitatory population receives external input and projects to a read-out layer in a purely feedforward manner. **B**. Sketch of the three plasticity rules: Spike-timing dependent plasticity (STDP) acts as a coincidence detector, strengthening (weakening) the synapse upon causality (anti-causality) of pre- and postsynaptic activity. Synaptic normalization (SN) normalizes all incoming synapses of a neuron (each represented by a distinct color) such that they sum up to unity: first, synaptic growth (or shrinking) due to STDP from step *t* to an intermediate step (*t* + 1)^*^ leads to the sum being bigger (or smaller) than unity. Second, synaptic competition for resources multiplicatively renormalizes all synapses at step (*t* + 1) such that they sum up to unity again. Intrinsic plasticity (IP) adapts the thresholds of the excitatory neurons to achieve a target firing rate *r*^*^. **C**. Sketch of the simulation procedure: during development, the excitatory population receives either noise or sequence input ***u***(*t*) and is subject to the plasticity rules. To quantify memory performance, plasticity is frozen and the excitatory population receives sequence input in both cases. A readout layer is trained by linear regression to predict the next sequence element ***u***(*t* + 1) from the current excitatory activity ***x***(*t*).

Here, the ***T***^***E***^ and ***T***^***I***^ are the thresholds, Θ(·) is the Heaviside step function and *W*_*ij*_ is the connection strength from neuron *j* to neuron *i*. Only a subset of excitatory neurons receives input ***u***(*t*) at each time step (see below). Note that effectively the neurons are fully leaky, i.e. only input at the timestep *t* contributes; as a result, all memory properties originate from the activity propagating via the synaptic connections *W*_*ij*_. All synapses from excitatory to inhibitory populations and vice versa (*W*^*IE*^ and *W*^*EI*^) are assumed to be present (supplying a “blanket” of dense inhibition, Karnani et al. [2014]). Inhibitory-inhibitory connections are considered absent (*W*^*II*^ = 0). The excitatory-excitatory connections (*W*^*EE*^) are sparse, and are subject to plasticity. This sparsity is random, with mean *λ* = 5 incoming and outgoing synapses per neuron. This gives a mean connection probability of 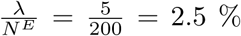. The sequence memory mechanism we propose shows qualitatively the same development in denser networks 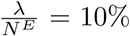; Fig. 3-S5).

### Plasticity rules

We model network development through the actions of activity-dependent plasticity mechanisms (Fig. 1B). Spike-timing dependent plasticity (STDP) acting on the excitatory-excitatory synapses *W*^*EE*^ is central to sequence learning. Two homeostatic mechanisms stabilize the network: synaptic normalization (SN) scales every neuron’s incoming synaptic weights in a multiplicative manner, such that they sum up to unity. Intrinsic plasticity (IP) adapts each excitatory neuron’s threshold 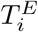 such that its activity matches a target rate of 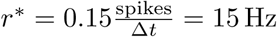.

### Input conditions

We study the development of the network (Fig. 1C) under two input conditions (Fig. 2A): “sequences” refers to structured input with temporal correlations, e.g. from sensory systems [Lazar et al., 2009, Klos et al., 2018], while “noise” refers to unstructured input, e.g. spontaneous cortical activity in a developmental stage before eye opening [Richter and Gjorgjieva, 2017, Triplett et al., 2018]. The “sequence” input [Lazar et al., 2009] is randomly chosen from a set of two sequences {*s*_1_, *s*_2_} of identical length (*L* = 10 steps) and structure: *s*_1_ = [*a*, (*L* − 2) × *b, c*]; *s*_1_ = [*d*, (*L* − 2) × *e, f*]. Each letter *l* activates a distinct set of 10 excitatory neurons, thus the mean input rate 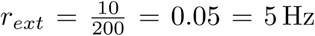 is constant. We construct the “noise” input by letter-like activation of 10 random neurons at each timestep, hence it has exactly the same rate as the sequence input but lacks any spatiotemporal correlations.

**Figure 2.**
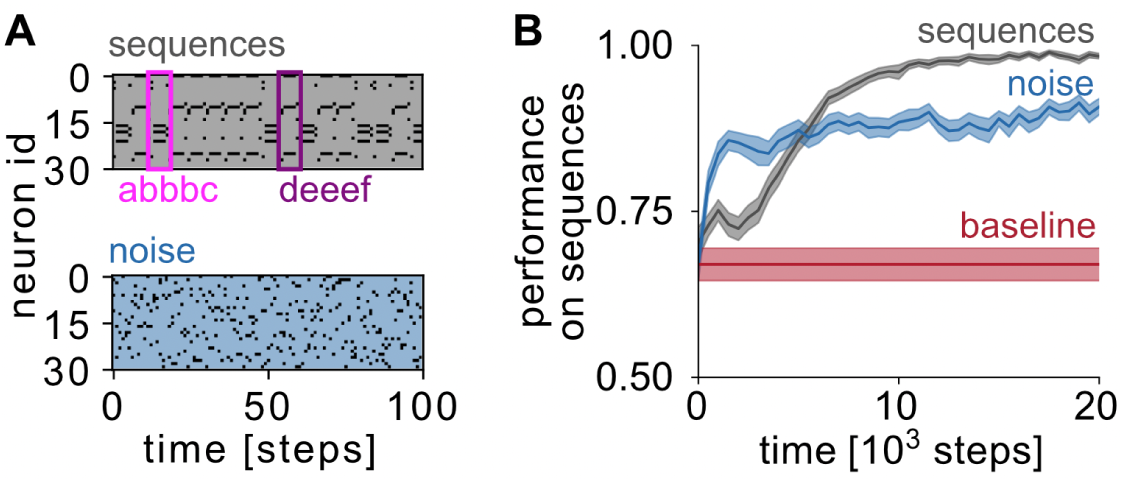
Development under noise input increases sequence memory performance. **A**. Development is simulated under two different input conditions: sequences model sensory input, noise models spontaneous activity. Shown are spike-rasters of input for 30 out of 200 neurons, the sequences *s*_1_ = [*a, b, b, b, c*] and *s*_2_ = [*d, e, e, e, f*] (see Methods) are marked in magenta and violet, respectively. **B**. Under noise input, sequence memory performance improves faster than under sequence input, but does not reach the maximum. Baseline refers to the performance of the random initial network. Lines: mean over *N* = 50 independent realizations; shaded areas: 33 − 66 % confidence interval thereof.

### Quantifying performance

The ability of the network to reliably represent sequences is always quantified using sequence input, independent of the input during development. This ability is very general, and can be used e.g. to *memorize* past stimuli and their order, as well as to *predict* future input. In detail, to quantify the network’s sequence memory performance, we freeze the network’s plasticity at the chosen time of development (Fig. 1C). We then use a linear readout layer to predict the next sequence element from the network state elicited by the previous element. For comparability, we used the same task and parameters as in the original publication [Lazar et al., 2009]. Here, optimal performance requires memory of the input that was presented 9 = *L* − 1 steps back.

## Results

We simulated the development of a recurrent neuronal network governed by activity-dependent plasticity rules [Lazar et al., 2009]. To quantify the network’s sequence memory performance, we freeze the network’s plasticity rules and use a linear classifier to predict the next step of a pre-defined sequence. First, we reproduced the known results for the SORN [Lazar et al., 2009]. As expected, already the random initialization represents a fairly powerful reservoir and already shows higher-than-chance sequence memory performance (*P*_0_ = 0.65 compared to *P*_*chance*_ = 0.008, mean across *N* = 50 realizations, Fig. 2B, red). Moreover, after about 10^4^ steps of development, network plasticity enables almost perfect performance (*P* = 0.98, mean across *N* = 50 realizations, Fig. 2B, gray) in memorising the sequences seen during development.

### Development under noise input increases unspecific sequence memory performance and speeds up consecutive learning of specific sequences

We proceeded to investigate whether the specific sequences are necessary as input during development, or whether development under unstructured, spontaneous activity would already be sufficient to increase sequence memory performance. We modeled spontaneous activity from other areas as binary noise input with the same rate as the sequence input (Fig. 2A). Interestingly, the development under noise input alone increases the memory performance on the sequences significantly, even surprisingly quickly. Already within 10^3^ timesteps (≈100 sequences), performance increased to *P* = 0.85 *> P*_0_ (Fig. 2B, blue). However, performance then saturated and never reached that of development under the sequence input (*P* = 0.90 vs *P* = 0.98). The increase in sequence memory after development under noise is even greater in larger and denser networks (Fig. 2-S1). It is quite surprising that the network improves its performance on sequence prediction more quickly, if it developed *not* under those sequences, but under noise instead. This advantage is maintained for a considerable period. Only after 5000 timesteps, the “sequences” condition outperforms the “noise” condition. This result suggests that networks **pre-shaped under noise** could learn sequences faster.

To test whether pre-shaping under noise boosts subsequent sequence learning, we designed a two-phased development process: first the network develops under noise input for a pre-shaping period of *T*_*pre*_ steps, afterwards we switch to sequence input (*t* = 0 in Fig. 3A). This transition could reflect a developmental transition from e.g. closed to open eyes. During the noise phase, the memory performance develops as described above (Fig. 2B). Upon onset of sequence input, two different cases emerge: Given long pre-shaping periods (*T*_*pre*_ *>* 5 ·10^3^), the performance grows quickly towards the maximum (Fig. 3A, dark blue). However, if pre-shaping periods were shorter than *T*_*pre*_ *<* 5 ·10^3^, performance shows an initial dip after sequence input before it proceeds to increase towards the maximum (Fig. 3A, light blue). Even with the dip, none of the pre-shaped networks was ever worse than the naive network (*T*_*pre*_ = 0, Fig. 3A, lightest blue).

**Figure 3.**
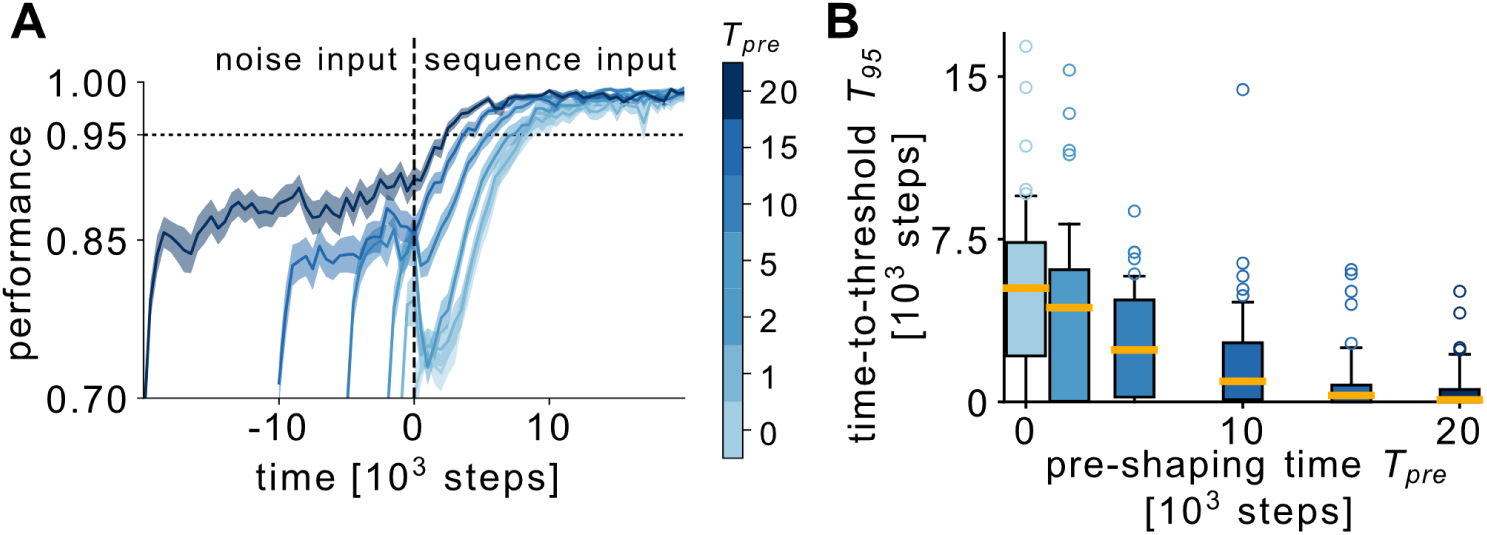
Pre-shaping under noise input speeds up subsequent learning of specific sequences. Two-phased development process with spike-timing dependent and homeostatic plasticity: First, development under noise input for different pre-shaping periods *T*_*pre*_. Then, development continues after onset of sequence input (*t* = 0, dashed vertical line in A). **A**. Pre-shaped networks (*T*_*pre*_ *>* 0) initially maintain higher sequence memory performance than those starting from random initial conditions (*T*_*pre*_ = 0) until all networks converge to similar performance. Lines: mean over *N* = 50 independent realizations; shaded areas: 33 − 66 % confidence interval thereof. **B**. The time *T*_95_ to reach 95 % performance decreases with longer pre-shaping periods *T*_*pre*_. Boxplot of the *T*_95_ distribution: boxes extend from 25th to 75th quantile, whiskers from 10th to 90th quantile. Thick bar inside the box denotes the median, colored circles show outliers. Distributions were estimated from *N* = 50 independent realizations for each pre-shaping period *T*_*pre*_.

To quantify how much the pre-shaping speeds up subsequent sequence learning, we introduce an arbitrary performance threshold of 95 % and measure the time *T*_95_ that each network required to reach that threshold: longer pre-shaping periods systematically shorten the time to reach that threshold (Fig.3 B). The speed-up of sequence learning after development under noise is even more pronounced in larger and denser networks (Fig. 3-S3). This clearly shows that simple noise input shapes the network in a way that speeds up subsequent sequence learning.

Would it then be wise to artificially prolong the noise pre-shaping phase (e.g. by *not* opening eyes immediately after birth or hatching)? To answer this question, we need to count the absolute development time (ranging [0, *T*_*pre*_ + *T*_95_]). On this time axis, networks with noise pre-shaping reach the performance threshold of 95% later than those that start with sequences (Fig. 3-S1A). Thus in absolute terms, the noise pre-shaping does not shorten the time to reach 95% performance. However, given any pre-shaping, a hatchling or newborn will always show better performance on sequences than without. Moreover, a sufficiently long pre-shaping period can even prevent the dip in performance after sequence onset. Thus, pre-shaping under noise is less efficient than learning from sequences when it comes to reaching maximal performance, however, sufficiently long pre-shaping strongly improves performance when the animal encounters the first structured sequences.

### IP and STDP+SN cause two distinct learning processes during the noise phase

Our results on pre-shaping under noise suggest the existence of two distinct developmental mechanisms in the network: fast structural changes under noise are beneficial for unspecific sequence memory (Fig. 3, light blue; Fig. 3-S2C,D) but do not facilitate consecutive learning of specific sequences (Fig. 3-S2C,D). On much longer timescales, structural changes emerge that facilitate consecutive sequence learning (Fig. 3, dark blue; Fig. 3-S2A,B). In the following, we uncover these structural changes and clarify the different roles of the underlying plasticity mechanisms (sketched in Fig. 4).

**Figure 4.**
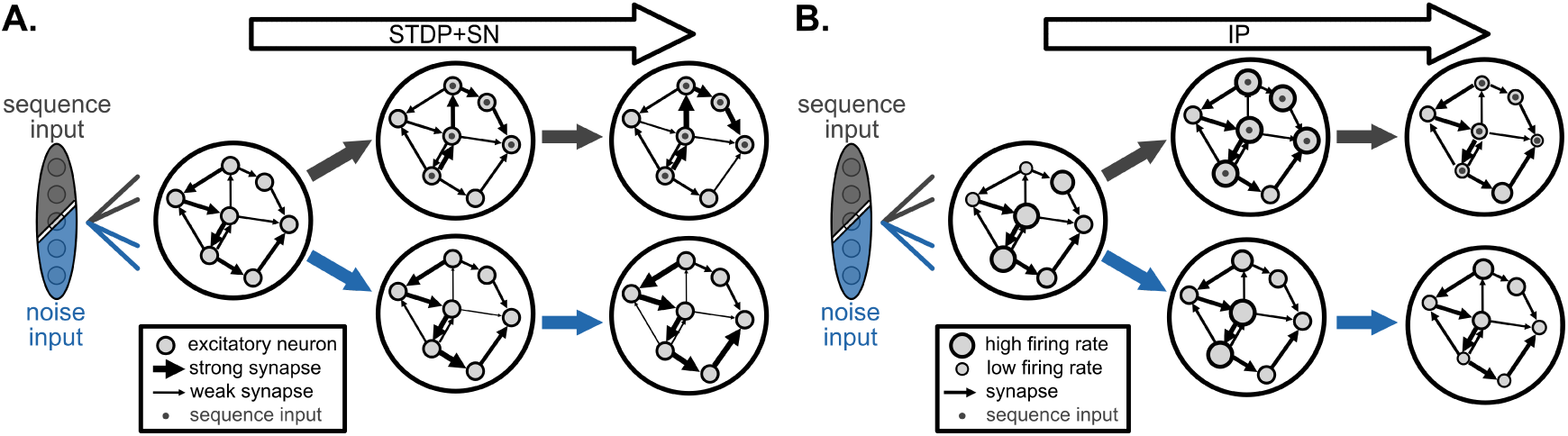
Spike-timing dependent plasticity and intrinsic plasticity have different effects on network structure. Cartoons of the effects of different input types and plasticity rules in isolation on network structure. **A**. Under sequence input, spike-timing dependent plasticity and synaptic normalisation (STDP+SN) imprint the input sequences into the network structure, regardless of initial synaptic weights. Thus finally, the preferential paths match the input sequences. Under noise input, STDP+SN amplifies those synapses which are strong in the initial synaptic matrix, strengthening and prolonging preferential paths that were already present. These paths can later be harnessed for learning of specific sequences. **B**. Intrinsic plasticity (IP) in isolation only alters the firing thresholds, leaving the synaptic weights unchanged. Under sequence input, IP quickly lowers the firing rate of neurons affected by the sequence input. Under noise input, IP first equalizes firing rates and later lowers the firing rates of neurons with strong incoming synapses. Firing rates are here “measured” under sequence input because those are relevant for sequence memory performance.

Already from considerations of effective timescales of the plasticity rules, we can pinpoint that IP is responsible for the rapid, unsustained performance increase, whereas STDP is responsible for the slow structural changes beneficial for subsequent sequence learning: Both timescales had been set [Lazar et al., 2009] to use the same **nominal learning rate** *η*_*IP*_ = *η*_*ST DP*_ = 10^−3^, but the **effective learning rates** differ: IP occurs at every timestep, independent of whether the neuron spiked or not, whereas the effective rate at which STDP occurs,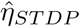, depends on the post- and presynaptic spikin activity. For uncorrelated post- and presynaptic spiking with target rate *r*^*^, STDP occurs with 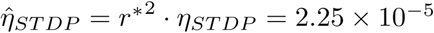, but even perfect correlation between spikes only gives 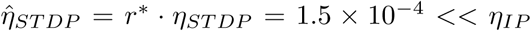, making STDP at least ten times slower than IP. Therefore, we expect IP to dominate the early development (up to 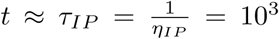) and the effects of STDP+SN to emerge later (after 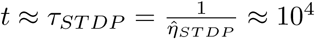).

To isolate the effects of different plasticity rules on sequence memory performance, we let networks develop under noise input but used either IP alone (updating the thresholds), or STDP+SN alone (updating the synaptic connections). To expose the characteristic timescales of network development, we use a logarithmic timeaxis in Figure 5 (note that the development slows down after *t >* 5 × 10^4^, but does not converge completely, see also Lazar et al. [2009]). Indeed, networks developing under IP alone do show the rapid rise in performance within *τ*_*IP*_ steps (Fig. 5A, pink). However, they cannot sustain this high performance for more than *τ*_*IP*_ steps (Fig. 5A), nor after sequence onset (Fig. 3-S2C,D). In contrast, networks developing under STDP+SN without IP lack this rapid rise, and instead show a slow climb in performance (starting after *t* = *τ*_*IP*_) up to the plateau observed in the STDP+SN+IP case (Fig. 5A, green). Additionally, they can sustain this plateau after sequence onset (Fig. 3-S2A,B). Concerted action of the three plasticity rules combines the best of two worlds and results in a rapid rise in performance that is sustained for all future times (Fig. 3A). Only the concerted action, too, gives rise to a network where all neurons participate with physiological firing rates and synaptic weights follow a long-tailed distribution (Fig. 5-S1, see also Lazar et al. [2009], Zheng et al. [2013]).

**Figure 5.**
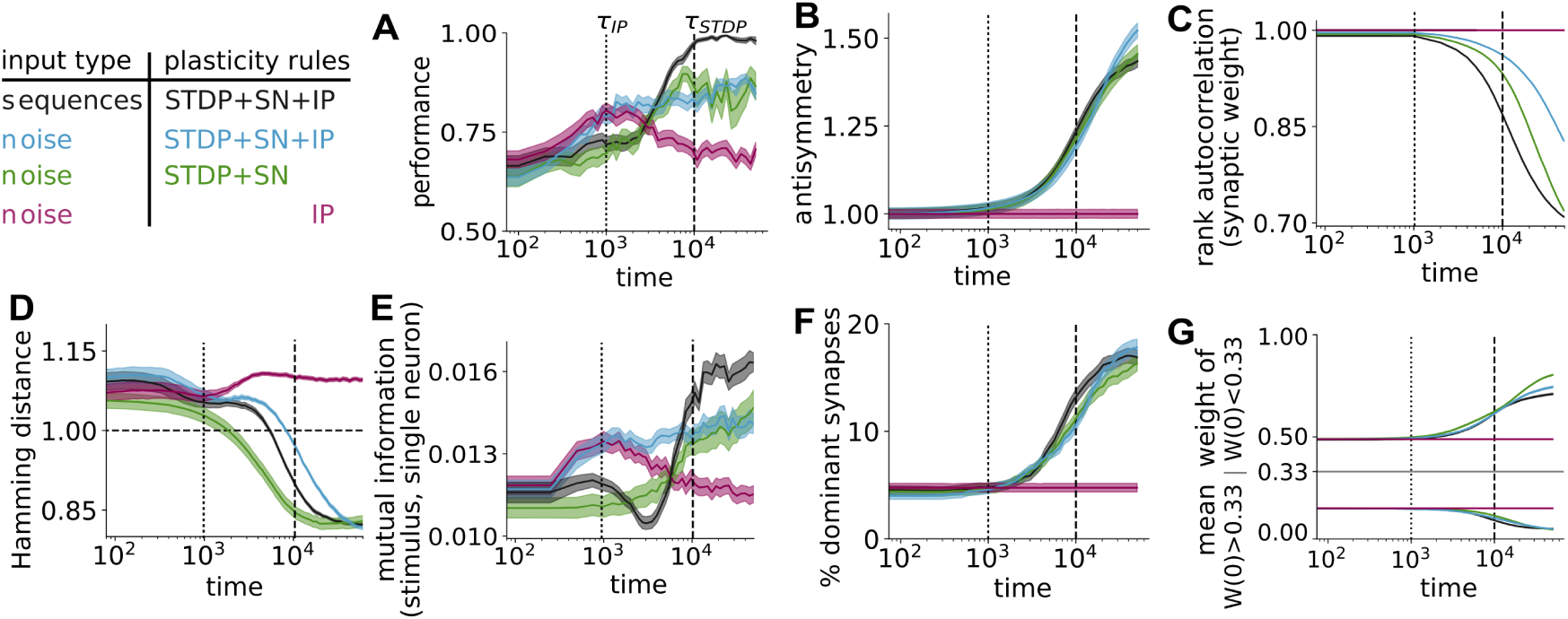
Spike-timing dependent plasticity and synaptic competition embed preferential paths into network structure. Isolated effects of the different plasticity rules on network development can be disentangled, because their characteristic time-scales differ (*τ*_*IP*_ ≈ 10^3^, *τ*_*STDP*_ ≈ 10^4^). **A**. Intrinsic plasticity (IP) under noise input (pink) boosts sequence memory performance quickly (peak at *t* = *τ*_*IP*_), but transiently (decayed at *t* = *τ*_*STDP*_). Spike-timing dependent plasticity and synaptic normalisation (STDP+SN) under both noise and sequence input cause a slow, but lasting increase in performance (start at *t* = *τ*_*IP*_, plateau at *t* = *τ*_*STDP*_). **B**. The antisymmetry of the synaptic matrix is increased by STDP+SN, but not by IP, independently of input type (noise or sequence). Its growth only starts after *t* = *τ*_*IP*_ and slows down after *t* = *τ*_*STDP*_ - paralleling the slow, lasting performance increase (panel A, green & black). The antisymmetry is normalized by the value at *t* = 0. **C**. Rank autocorrelation of the synaptic weights under noise input decays slower than under sequence input. Thus, synaptic weights at *t* = 0 largely determine those at *t* = *τ*_*STDP*_ under noise, while they are quickly forgotten under sequence input. **D**. The Hamming distance shows that networks with STPD+SN develop from chaotic, slightly supercritical to more stable, subcritical dynamics within *t* = *τ*_*STDP*_. **E**. The lagged mean mutual information between stimulus identity and single neuron activity shows similar transient peaks and plateaus as the performance (in A), indicating reliable propagation of input among neurons. **F**. The proportion of dominant synapses 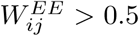 shows similar increase and plateau as the antisymmetry for both noise and sequence input. Along these dominant synapses, activity propagates reliably - they form the preferential paths that support sequence memory. **G**. The mean of initially weak synapses decreases further with development, while the mean of initially strong synapses grows, evidencing the positive self-feedback of STDP and synaptic competition. Lines: mean over *N* = 50 independent realizations; shaded areas: 33 − 66 % confidence interval thereof.

Because IP only transiently boosts performance (Fig. 5, pink; Fig. SR-1A) and lacks a positive pre-shaping effect (Fig. 3-S2C), we only summarise its structural effects and refer the interested reader to the supplementary results. The rapid rise in performance (Fig. 5A; Fig. SR-1A) is due to IP under noise input rapidly bringing the network activity average towards target rate (Fig. SR-1C), where performance of random networks is almost maximized (Fig. SR-1B). Prolonged development leads to IP slowly silencing neurons that are receiving excess recurrent input from strong incoming synapses (Fig. SR-1D,E; Fig. 5-S1D). Because such neurons are central to the reliable propagation of sequences (Fig. SR-1A,D,E; see also next section), IP acting in isolation is detrimental to sequence memory performance in the long run (Fig. 5A, pink).

### Preferential paths form the structural substrate of sequence memory

Because STDP+SN in isolation has a positive (Fig. 5A), sustained pre-shaping effect on performance (Fig. 3-S2A,B), we will now analyse its structural effects in detail. The STDP rule naturally increases one synaptic direction at the expense of the reverse direction, leading to a natural increase in antisymmetry [Klos et al., 2018]. Indeed, the antisymmetry of the weight matrix *W*^*EE*^ as measured by the Frobenius norm ∥*W*^*EE*^ − (*W*^*EE*^)^*†*^∥ _*ℱ*_ only starts to increase after *τ*_*IP*_ = 10^3^ and shows its strongest increase after about 10^4^ steps (Fig. 5B). This corresponds to the effective timescale of STDP, *τ*_*ST DP*_ = 10^4^, and parallels the performance increase in the STDP+SN condition. Hence, the lasting increase in performance originates from STDP+SN increasing the antisymmetry of the network.

How does this increase in antisymmetry help the network to memorize sequences? Along the “strong” directions, activity can propagate with high reliability regardless of the background activity in the network - activity thus propagates along **preferential paths** [Fiete et al., 2010, Klampfl and Maass, 2013, Hemberger et al., 2019]. The propagation probability along a path increases with the synaptic strength in relation to the thresholds. Interestingly, the share of “dominant” synapses - those which are stronger than all other incoming synapses combined (i.e. 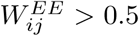) - shows a very similar development to the antisymmetry (Fig. 5F). Such dominant synapses have been observed in vivo [Song et al., 2005, Lefort et al., 2009, Hemberger et al., 2019] and are thought to play an important part in network function [Lefort et al., 2009]. Taken together, these dominant synapses establish preferential paths along which activity can propagate reliably for multiple timesteps. This increased reliability is directly evidenced by the parallel rise of the delayed mutual information (a non-linear generalization of correlation, [Cramer et al., 2019]), measured between sequence identity and single neuron activity (Fig.5E). Hence, the rise in sequence memory performance seen in Fig. 5A relies on STDP imprinting preferential paths into the network structure.

These preferential paths form under both sequence and noise input (Fig. 5F). They are already present in the initial random connectivity matrix, but short and few (Fig. 5F; see also 6A,B,C). Under noise input, STDP+SN work together to elongate and strengthen those present (Fig. 5G). Under sequence input, STDP+SN establish preferential paths starting at the input neurons, with much less dependency on initial synaptic weight (Fig. 5C). Hence, those paths formed under noise input do enable memory of specific sequences but are sub-optimal compared to those that form under sequence input (Fig. 5A). However, due to the positive self-feedback of STDP+SN those formed under noise input are strengthened and prolonged towards optimality under consecutive sequence input (Fig. 3A), facilitating fast learning of specific sequences (Fig. 3B).

### Formation of preferential paths gives rise to subcritical dynamics

Contrary to the long-standing hypothesis that the information processing capability of recurrent neural networks is optimal when their dynamics are critical [Beggs, 2008, Boedecker et al., 2012], we find that development increases performance while driving the dynamics from slightly supercritical to subcritical. We use the Hamming distance *D*, a one-step estimation of the network’s effective Lyapunov exponent, as a measure of criticality [Lazar et al., 2009]: *D >* 1 marks the regime of supercritical dynamics; *D* = 1 marks the critical point, and *D <* 1 marks subcritical dynamics. At the onset of development, the network operates in a chaotic, slightly supercritical (*D* ≈ 1.05 − 1.10, Fig. 5D) or “asynchronous-irregular” regime of spiking dynamics [Brunel, 2000] that reproduces key features of single-neuron cortical dynamics [Hartmann et al., 2015]. Under both noise and sequence input, it develops towards a subcritical (*D* ≈ 0.85) or “reverberating” regime that characterizes large-scale cortical dynamics *in vivo* [Wilting and Priesemann, 2019] and can be maintained by homeostatic mechanisms [Zierenberg et al., 2018, Ma et al., 2019]. The timescale of this **plasticity-driven decrease in criticality** matches both that of (i) the formation of preferential paths (Fig. 5G) and (ii) the STDP+SN-driven increase in mutual information between single-neuron activity and stimulus (Fig. 5E). This indicates that the preferential paths formed by STDP+SN are not only strong enough to reliably propagate memory of the stimulus despite the recurrent background activity (Fig. 5E), but also dampen external perturbations (Fig. 5D). Crucially, criticality is decreasing while the performance is increasing 5A,D). This adds another example to the increasing body of evidence (Lazar et al. [2009], Del Papa et al. [2017], Cramer et al. [2019], reviewed in Wilting et al. [2018]) that plasticity rules can drive network dynamics away from criticality while increasing performance in a specific task.

### Spontaneous activity contains endogenous sequences propagating along preferential paths

The presence of preferential paths in the connectivity shapes the spontaneous activity of the network [Klampfl and Maass, 2013, Hartmann et al., 2015]. To uncover the relation between connectivity and activity, we first re-order the synaptic matrix *W*^*EE*^ (Fig. 6A) such that the preferential paths lie on the lower secondary diagonal. We achieve this by starting from a random neuron and defining its “follower” as the post-synaptic neuron of the largest outgoing synapse. If the so-defined “follower” is already present in the ordering, the post-synaptic neuron of next-largest outgoing synapse is selected. This method of sorting reveals multiple preferential paths of different length on the lower secondary diagonal (Fig. 6B) that grow during development (Fig. 6C). Yet, the off-diagonal entries of *W*^*EE*^ are still sparsely populated (Fig. 6A,B), preserving a physiological network structure.

**Figure 6.**
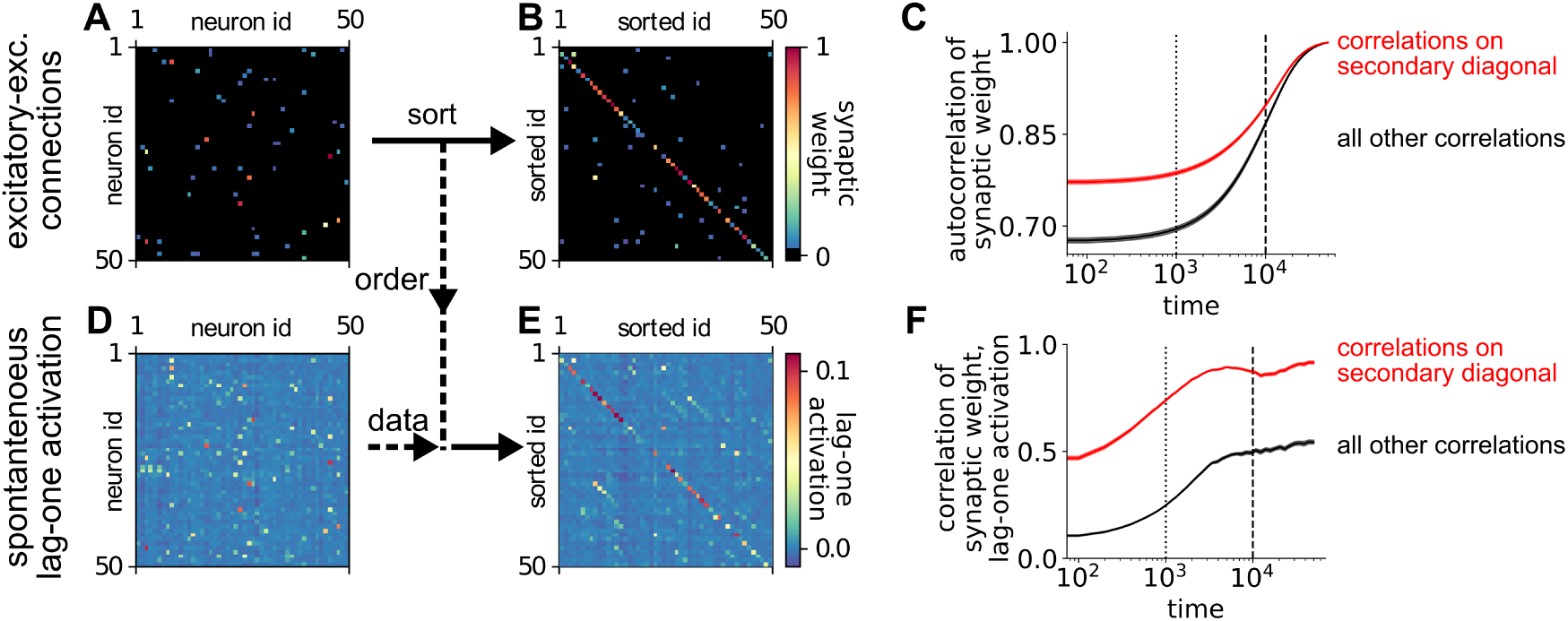
Spontaneous activity displays endogenous sequences propagating along preferential paths. The excitatory-excitatory connections *W* ^*EE*^ (panels A,B) shape spontaneous activity, here visualized as the lag-one activation matrix *Q*^*EE*^ (panels D,E). The panels show 50×50 subsets of the 200×200 matrices *W* ^*EE*^ and *Q*^*EE*^ after *T* = 10, 000 steps of development under noise input. Synaptic weights 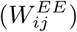 or lag-one activations 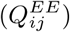 during spontaneous activity are in color, 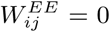 is depicted in black. **A**. Synaptic matrix *W* ^*EE*^ shown in neuron id space (unordered). **B**. Sorting *W* ^*EE*^ along strongest “follower” synapses starting from a random neuron (thereby defining sorted id space) reveals multiple disjoint preferential paths of strong synapses on the lower secondary diagonal. **C**. Autocorrelating the secondary diagonals of *W* ^*EE*^ in sorted ID space (applying the sorting from *W* ^*EE*^(50, 000) to every other timestep *W* ^*EE*^(*t*), red) shows that preferential paths present in the initial matrix predict preferential paths after development very well (red compared to black). **D**. Lag-one activation matrix *Q*^*EE*^ shown in neuron id space (unordered). **E**. Applying the sorting obtained from *W* ^*EE*^ to *Q*^*EE*^ reveals multiple disjoint endogenous sequences propagating along preferential paths (on the lower secondary diagonal). **F**. Correlating the lower secondary diagonals of *W* ^*EE*^ and *Q*^*EE*^ in sorted ID space (applying the sorting from *W* ^*EE*^(*t*) to *Q*^*EE*^ at every timestep, red) reveals that preferential paths predict endogenous sequences very well (red compared to black). Lines show the mean of Pearsons’ correlation coefficient over *N* = 50 independent realizations, shaded areas show the 33 − 66% confidence interval thereof.

To confirm that connectivity shapes spontaneous activity, we apply the ordering obtained from the synaptic matrix *W*^*EE*^ to the lag-one activation matrix *Q*^*EE*^ (Fig. 6D). The lag-one activation matrix 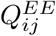 of neural activity *x*_*i*_(*t*), *x*_*j*_(*t* − 1) was estimated from a frozen network under noise input. Specifically, lag-one activation 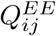 was defined as the mean of sequential activations, corrected for the individual firing rates: 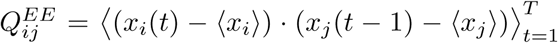, with *T* = 10^4^. Applying the ordering obtained from *W*^*EE*^ to *Q*^*EE*^ reveals **endogenous sequences** [Carrillo-Reid et al., 2015, Dechery and MacLean, 2017, Hemberger et al., 2019] on its lower secondary diagonal (Fig. 6E): multiple neural activations following each other with higher-than chance probability.

To quantify how well the presence of preferential paths in the connectivity predicts the presence of endogenous sequences in spontaneous activity, we correlated the entries of the follower-sorted matrices *W*^*EE*^ and *Q*^*EE*^. We find that the presence of preferential paths in *W*^*EE*^ predicts endogenous sequences in *Q*^*EE*^ well: correlating the secondary diagonals of *W*^*EE*^ and *Q*^*EE*^ yields *ρ*_*on*_ ≥ 0.5 even at *t* = 0 (Fig. 6F, red), much bigger than the correlation of off-diagonal elements (*ρ*_*off*_ (*t* = 0) ≤ 0.1, Fig. 6F, black). This remains true during development and upon plateauing: *ρ*_*on*_(*t* = 10, 000) ≈ 0.85 *> ρ*_*off*_ (*t* = 10, 000) ≈ 0.5 (Fig. 6F). The plateau is already reached at a time intermediate between the timescales of IP and STDP (*η*_*IP*_ = 10^3^ ≤ *t* = 5 ·10^3^ ≤ 10^4^ = *η*_*ST DP*_). This evidences that endogenous sequences arise from structures created by STDP+SN but do not need them to be fully developed. Thus, already short development under unstructured input gives rise to endogenous sequences present in spontaneous activity.

## Discussion

We addressed whether spontaneous activity prior to onset of sensory input can prepare the cortex for memorising input sequences. We analysed the development of a recurrent neural network model subject to activity-dependent plasticity rules. Integrating STDP, synaptic normalisation and a rate homeostasis, we demonstrate for the first time that development under unstructured spontaneous activity gives rise to improved sequence memory while maintaining a biologically plausible connectivity and firing rate regime. We identified the origin of this improved sequence memory: preferential paths in the synaptic matrix, which are reflected in the spontaneous activity as endogenous sequences. Finally, we highlight that this emerging network structure benefits fast learning of specific sequences, and may improve “performance” of a newborn after hatching.

### The SORN model reproduces key features of cortical dynamics

Any neural network model can incorporate some of the properties of real neural systems, but still has to reduce the biological complexity considerably. Our model, for example, uses a fairly simplistic spiking mechanism (binary threshold neurons), but still reproduces key properties of cortical spiking behaviour. In cortex, spiking neurons receive loosely balanced excitatory and inhibitory input through strong synapses (current/threshold ratio on the order of 10) [Sanzeni et al., 2020]. They might operate in a chaotic regime [Brunel, 2000] characterised by a large variance of activity [London et al., 2010] and exponential distributions of inter-spike intervals [Goris et al., 2014]. Our model uses binary threshold neurons that receive loosely balanced input through synapse of moderate strength (current/threshold ratio on the order of 3, Fig. 5-S1C). Yet, it also operates in a chaotic regime prior to learning (Fig. 5D) and has been shown to reproduce key features of cortical spiking [Hartmann et al., 2015, Del Papa et al., 2017]: high variability during rest (exponential ISI distributions, Fano factor close to 1) that is quenched upon stimulus onset. Last, a recent generalization of our model to leaky integrate-and-fire neurons [Miner and Triesch, 2016] reproduced virtually the same key features of cortical spiking behaviour.

To leverage transient states for computation, neuronal networks cannot be fully chaotic, but need some spatiotemporal structure in their activity [Buonomano and Maass, 2009, Boedecker et al., 2012]. While randomly connected balanced networks can give rise to multiple regimes of chaotic activity [Mastrogiuseppe and Ostojic, 2017] that are thought to reproduce single-neuron spiking statistics [London et al., 2010], their chaos needs to be tamed to explain large-scale spiking statistics [Tomm et al., 2014, Wilting and Priesemann, 2019] and support transient computation [Lazar et al., 2009, Laje and Buonomano, 2013]. This can be achieved through the use of non-random connectivity [Tomm et al., 2014, Landau and Sompolinsky, 2018, Spreizer et al., 2019], hidden low-rank matrices [Bondanelli and Ostojic, 2020], or unsupervised learning mechanisms [Del Papa et al., 2017, Zierenberg et al., 2018]. Interestingly, two studies showed that even supervised learning of functional transient computations quenches chaos, in essence reinforcing the intrinsic transient dynamics of the network [Laje and Buonomano, 2013]. In our model, intrinsic transient network dynamics and local plasticity mechanisms form functionally relevant structures that are embedded into close-to-chaotic network activity.

### Towards more detailed models of sequence memory development

We studied the development of sequence memory, which relies heavily on the plasticity of excitatory-excitatory connections [Fiete et al., 2010]. We thus focused on the plasticity of excitatory neurons, and neglect connections between inhibitory neurons altogether. The richness of the physiology and function of inhibitory-inhibitory connections are only beginning to be unraveled [Jiang et al., 2015]. By now is clear that dense inhibitory circuits with broad tuning supply an unspecific “blanket” of inhibition to the excitatory population [Fino and Yuste, 2011]. Connections between inhibitory neurons are not deemed necessary for this blanket [Isaacson and Scanziani, 2011], but have been implicated in maintaining gamma rhythms and precise spike timings [Isaacson and Scanziani, 2011] - which are out of the scope of our discrete-time network simulation. Yet, recent models that use leaky integrate-and-fire neurons, operate in continuous time and incorporate inhibitory-inhibitory connections exhibit the same structural substrates of sequence memory we observed: increase of synaptic strengths along the sequence direction, and spontaneous replay of the learned sequence [Klos et al., 2018]. A complementary study of similar biological detail showed that endogenous sequences can arise from the interaction of STDP and spontaneous activity [Izhikevich et al., 2004]. Since the two main ingredients of our proposed sequence memory mechanism exist in these more detailed models, we are confident that our proposed mechanism also applies to spiking networks with more detailed connectivity.

Previous studies have analysed the development of mainly excitatory networks with spike-timing dependent plasticity under spontaneous activity, and some speculated on its impact on sequence memory: Analytical studies showed that STDP can autonomously develop feed-forward structures [Ocker et al., 2015, Ravid Tannenbaum and Burak, 2016, Montangie et al., 2020]. Numerical studies confirm this result: Networks equipped with STDP and a form of synaptic normalisation converge to form multiple synfire chains under both sequence and noise input [Fiete et al., 2010, Weissenberger et al., 2017]. Due to the lack of a second homeostatic mechanism [Zenke and Gerstner, 2016] in that network, however, all of the non-feedforward synapses die and the rich recurrent connectivity found *in vivo* is not well captured [Fiete et al., 2010]. It was speculated that such structures might facilitate one-shot-learning [Weissenberger et al., 2017]. Moreover, unsupervised pre-shaping by STDP in excitation-only networks can speed-up *supervised* sequence learning [Melchior et al., 2019]. Our study now extends theories of autonomous chain formation to networks in various aspects: physiologically, we incorporate *an inhibitory population and multiple homeostatic mechanisms*, and functionally, we are the first to demonstrate (i) that development under noise improves sequence memory and (ii) speeds up *unsupervised* sequence learning.

We considered two different forms of input during development in our *in silico* study: spatiotemporally correlated input as a model of sensory evoked activity, and uncorrelated input as a first minimal model of spontaneous activity. *In vivo*, spontaneous cortical activity contains some spatiotemporal correlations. The exact nature of these correlations varies between developmental stages: early on, bursts of poorly correlated cortical activity are driven by peripheral input [Colonnese et al., 2017]. Shortly before eye opening, subnetworks in visual cortex are active independent of peripheral drive [Pietri et al., 2017]. While structured input from the periphery is required for the complete maturation of cortical circuitry [Ko et al., 2014], its absence does not prevent the emergence of specific functional recurrent connectivity [Ko et al., 2014, Pietri et al., 2017]. We thus chose to study the impact of uncorrelated input, and view the emerging structures as a lower bound of structure that will be generated by spontaneous cortical activity. We found that both correlated and uncorrelated input generated self-amplifying preferential paths - they differed quantitatively, not qualitatively. Hence, it is to expect that STDP and homeostatic plasticity under spontaneous activity will shape recurrent networks to contain some form of preferential paths, regardless of the exact statistics of the spontaneous activity.

### Preferential paths as building blocks of sequence memory

We find that preferential paths in the network structure are correlated with sequences in the spontaneous activity of the network dynamics, reminiscent of endogenous sequences observed *in vivo* [Carrillo-Reid et al., 2015, Dechery and MacLean, 2017]. The preferential paths in our network are built by strong synapses, which are over-represented *in vivo* [Song et al., 2005, Lefort et al., 2009] and thought to be crucial for cortical computation [Lefort et al., 2009]. Such strong synapses allow the artificial stimulation of a single neuron to reliably cause sequential activation of neural assemblies [Hemberger et al., 2019] and might also underlie endogenous sequences *in vivo* [Hemberger et al., 2019]. Similar to endogenous sequences outlining the realm of activity evoked by sensory stimuli *in vivo* [Luczak et al., 2009, Carrillo-Reid et al., 2015], the preferential paths formed in our network facilitate reliable propagation of both externally and internally generated sequences.

The development of preferential paths in the absence of structured input in our *in silico* model offers an explanation for the emergence of endogenous sequences observed in the *in vivo* spontaneous activity [Luczak et al., 2007, Dechery and MacLean, 2017]. Initially, the plasticity rules acting on unstructured input amplify asymmetries already present in the network structure, creating preferential paths. As the paths strengthen, they are activated more and more, thereby shaping the spontaneous activity to display sequences. Thus, initially unstructured spontaneous activity shapes the connections such that endogenous sequences are spontaneously activated. These sequences could then be recruited by sensory stimuli [Carrillo-Reid et al., 2015] and the synapses connecting them strengthened to form a memory trace after only a few reactivations.

Because synaptic plasticity in neocortex and hippocampus is similar [Cai et al., 2007], this mechanism might even be extended to cover a similar phenomenon observed in hippocampus: the “pre-play” of neural sequences during rest that will later code for memories acquired in a navigation task [Dragoi and Tonegawa, 2011]. Indeed, a recent modelling study has demonstrated that pre-played endogenous sequences can act as a template of rapid memory formations [Haga and Fukai, 2018], but did not show how they may arise. We here suggest that endogenous sequences emerging from the interplay of spontaneous activity and plasticity could underlie hippocampal pre-play.

## Conclusion

Using an established neural network model with plasticity [Lazar et al., 2009, Hartmann et al., 2015, Wang et al., 2017], we showed that sequence memory can develop in the absence of structured input. This result highlights the functional role of spontaneous activity during development. It also offers an explanation for endogenous cortical sequences recently observed *in vivo* [Carrillo-Reid et al., 2015, Dechery and MacLean, 2017, Hemberger et al., 2019] as well as the “pre-play” phenomenon observed in hippocampus [Dragoi and Tonegawa, 2011]. Furthermore, we provide a mechanistic understanding of how “pre-shaping” with unsupervised, local plasticity rules can greatly enhance the learning speed of artificial neural networks. If verified in experiment, the mechanism we propose could change the theory of memory: memory traces are not printed on a *tabula rasa*, but can be established quickly by harnessing building blocks already present in the brain.

## Methods

### Recurrent network model

We use the self-organized-recurrent neural network model (SORN, Lazar et al. [2009]) with the same parameters as in the original publication (table 1). The network has *N*^*E*^ = 200 excitatory and *N*^*I*^ = 0.2 × *N*^*E*^ = 40 inhibitory binary threshold neurons, respectively. The sequence memory mechanism we propose shows the same development in denser networks (*N*^*E*^ = 1000; Fig. 2-S1A, Fig. 3-S3A). Each neuron *i* sums the input it receives at time *t* = *t*_*k*_ linearly and spikes (*x*_*i*_(*t*_*k*_) = 1) if the sum exceeds its threshold *T*_*i*_, or else stays silent (*x*_*i*_(*t*_*k*_) = 0). The neurons are connected by weighted synapses, where *W*_*ij*_ is the connection strength from neuron *j* to neuron *i* (no self connections, *W*_*ii*_ = 0 ∀*i*). We simulate the network in discrete timesteps and interpret a single timestep as Δ*t* = 10 ms:

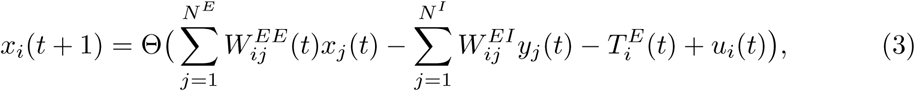

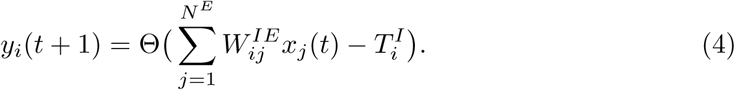

**Table 1.**
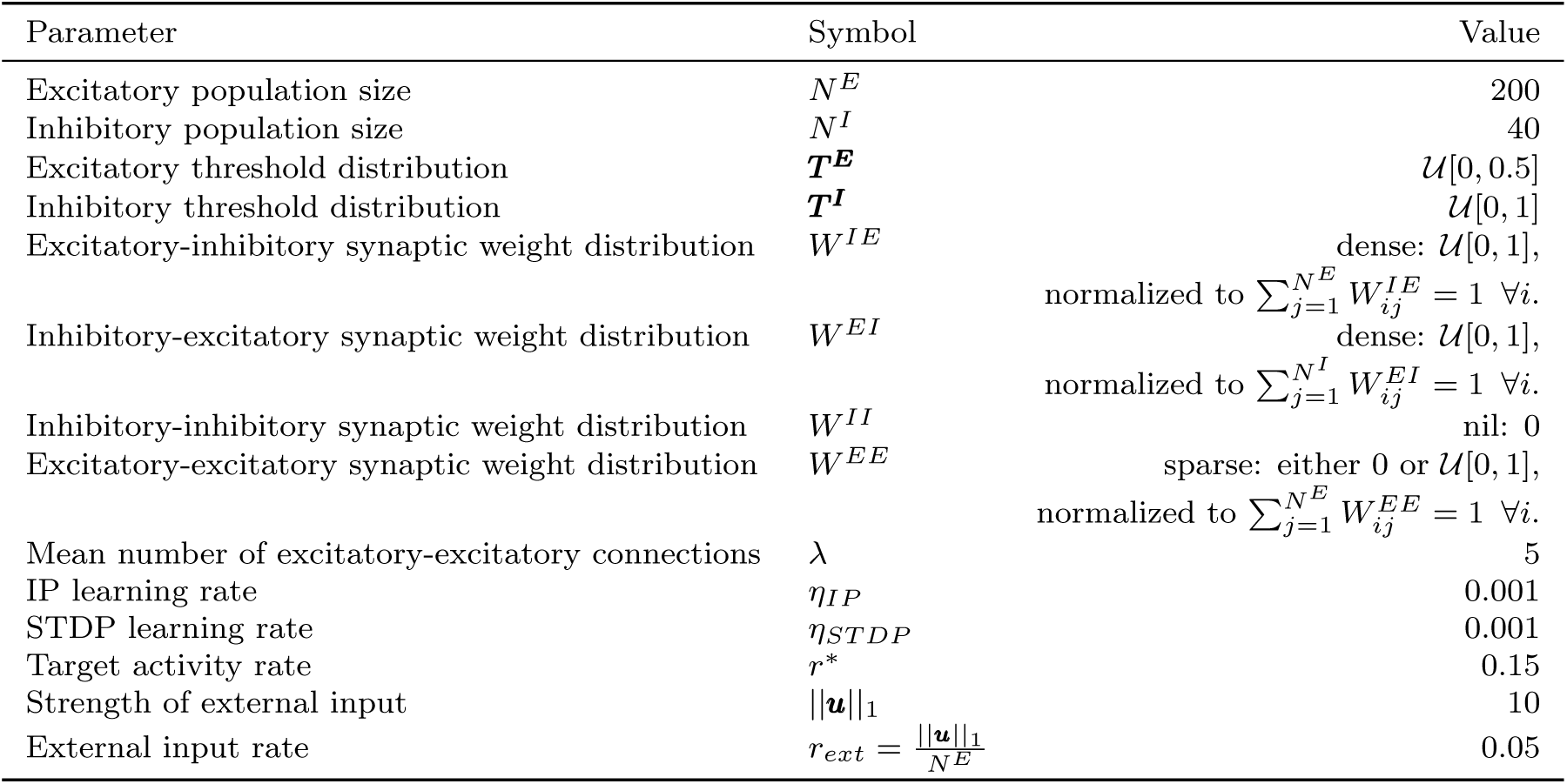
Initial Parameters.

Here, the Heaviside threshold function Θ(·) ensures that neural activity is a binary variable 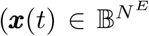 and 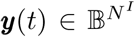, respectively). The external input (see below) 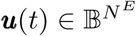 is the only source of stochasticity, introducing either high spatiotemporal correlations (sequence input) or no correlations (noise input).

Initially, the excitatory and inhibitory thresholds 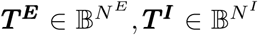 are drawn from the uniform distributions 𝒰[0, 0.5] and 𝒰 [0, 1], respectively. All synapses between excitatory and inhibitory neurons are present, thus 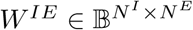 and 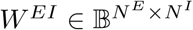 are dense matrices. In contrast, each excitatory-excitatory synapse is present with a probability of 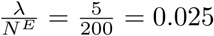, thus 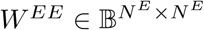 is a sparse matrix. The sequence memory mechanism we propose shows the same development in denser networks (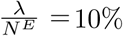; Fig. 2-S1B, Fig. 3-S3B). All inhibitory-inhibitory connections *W*^*II*^ are assumed to be nil. All synaptic weights are *W*_*ij*_ drawn from the uniform distribution 𝒰 [0, 1] and subsequently normalized such that the incoming connections to each neuron sum up to 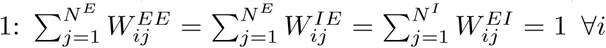. If all neurons spike with the same rate *r*^*^, then this normalisation assures that all neurons receive similar mean drive. The resulting distribution of weights 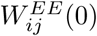 is shown in Figure 5-S1L. We found that denser networks need to be initialized with a log-normal distribution (*µ* = 0, *σ* = 1) prior to normalisation to generate the large variation of synaptic weights found *in vivo* [Song et al., 2005], and thus initialised all networks with *λ >* 5 in the “density scaling” plots (Fig. 2-S1B, Fig. 3-S3B) in this manner.

### Plasticity rules

Network development is simulated as the action of three distinct plasticity rules that change the connections between excitatory neurons *W*^*EE*^ and the threshold of excitatory neurons ***T***^***E***^ (see also Table 2 and Figure 1B).

**Table 2.** Variables changing during development.

| Variable | Symbol | Def | Value |
| --- | --- | --- | --- |
| Excitatory activity | $\mathbf{x}$ | 3 | propagates via $W^{EE}$, $W^{IE}$ |
| Inhibitory activity | $\mathbf{y}$ | 4 | propagates via $W^{EI}$ |
| Excitatory thresholds | $\mathbf{T}^E$ | 7 | subject to IP |
| Excitatory-excitatory connections | $W^{EE}$ | 5, 6 | subject to STDP and SN |
| propagation delay | $\Delta t$ | 3, 4 | 1 |
| integration time | $\Delta t$ | 3, 4 | 1 |
| effective IP timescale | $\tau_{STDP}$ | 7 | fast: 1,000 |
| effective STDP timescale | $\tau_{STDP}$ | 5 | slow: 10,000 – 100,000 |
| effective SN timescale | $\tau_{SN}$ | 6 | instant: 1 |

#### Spike-timing Dependent Plasticity

Most important for sequence learning, spike-timing dependent plasticity (STDP) strengthens connections between neurons that exhibit “causal” spiking activity. We use a form of STDP that detects causality on the timescale of a single timestep. In detail, an excitatory-excitatory synapse’s weight 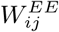 is only raised or lowered by a small amount *η*_*ST DP*_ if single-step causality or anti-causality is detected:

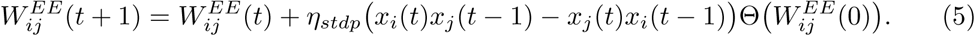

In the above definition, Heaviside’s step function 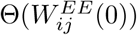 ensures that the plasticity rule only acts on connections present at *t* = 0, i.e. creates no new connections. Note that STDP acting in isolate can grow the synaptic weight towards infinity or shrink it towards zero, potentially destabilizing the network. To distribute the activity in the network evenly and counteract the destabilizing nature of STDP, we make use of additional homeostatic mechanisms that are sensitive to the total synaptic input and the post-synaptic firing rate.

#### Synaptic Normalisation

Synaptic normalisation (SN) is motivated by the competition of synapses for limited resources, keeping the sum of synaptic weights constant [Triesch et al., 2018]. SN multiplicatively rescales the values of incoming excitatory connections to a neuron such that they sum up to unity:

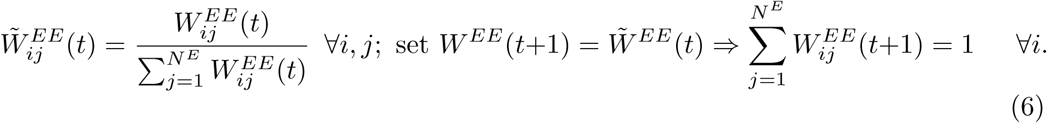

This rule acts instantly after synaptic weights are changed by STDP. Note that this rule does not change the relative strengths of synapses as established by STDP but regulates the total incoming drive a neuron receives, acting as a homeostatic mechanism.

#### Intrinsic Plasticity

Intrinsic plasticity (IP) regulates the firing rate of excitatory neurons based on their past firing activity. IP adapts each excitatory neuron’s threshold 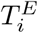 such that a global target firing rate *r*^*^ is met. At every timestep, the threshold is lowered by *η*_*IP*_ · *r*^*^ if the neuron just fired or raised by *η*_*IP*_ · (1 − *r*^*^) if it stayed silent:

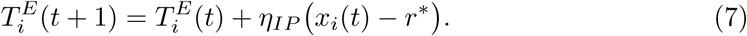

The target rate is 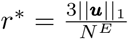, hence the number of input spikes is approximately one third the total number of spikes. The target rate *r*^*^ is identical for all neurons, thus IP spreads activity evenly in the network.

### Input conditions

We study the development of the network (Fig. 1C) under two input conditions (Fig. 2A): “sequences” refers to structured input with temporal correlations, e.g. from sensory systems [Lazar et al., 2009, Klos et al., 2018], while “noise” refers to unstructured input, e.g. spontaneous cortical activity in a developmental stage before eye opening [Richter and Gjorgjieva, 2017, Triplett et al., 2018].

#### Sequence input

The two sequences *s*_1_, *s*_2_ have identical structure and length *L* = 10 [Lazar et al., 2009], but consist of distinct sequence elements or “letters” *l* ∈ {*a, b, c*; *d, e, f*}:

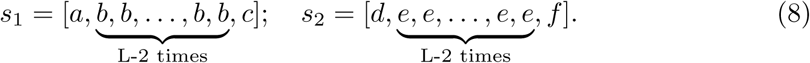

Each letter *l* activates a distinct set of 15 external neurons trough the corresponding input vector 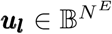, hence the mean external input rate is 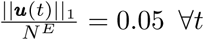. Note that this external input is far from uniformly distributed: all neurons not directly activated by presentation of a letter do not receive input, and those activated by the letters “b” or “f” receive an input rate that is higher than the target firing rate. The input vectors ***u***_***l***_ are chosen pseudo-randomly such that there is no overlap and are kept constant over time. The sequences are presented one letter at a time, and repeated in a random fashion. Thus, the “sequence” input condition introduces strong spatiotemporal correlations into the network dynamics.

#### Noise input

We designed a letter-like “noise” input that has exactly the same strength as the “sequence” input but lacks any spatiotemporal correlations. Specifically, 10 excitatory neurons are activated at every timestep, chosen in a random fashion. The probability to activate a given neuron *i* is uniformly distributed, hence the external input to the network has the same mean strength as in the “sequence” input condition, but lacks spatial structure and temporal correlations.

### Assessing sequence memory performance

To assess sequence memory performance *P*, we use the same counting task as described in Lazar et al. [2009]. In this task, the excitatory population receives a stream of external input *U* consisting of randomly interleaved presentations of two sequences *s*_1_ or *s*_2_. After this training phase, a readout layer *A* is trained via linear regression to predict the next sequence element *v*_*l*_(*t* + 1) from the current excitatory population activity ***x***(*t*). Here, the vector 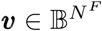 will be called the stimulus identifier, its dimensionality *N*^*F*^ = 6 is the number of letters (or “features”: *l* ∈ {*a, b, c*; *d, e, f*}, appearance of *l*: *v*_*l*_ = 1, absence of *l*: *v*_*l*_ = 0). The performance *P* of the readout layer *A* is afterwards assessed in a testing phase on an independently generated stream of external input *U′* consisting of the same sequences *s*_1_,*s*_2_, but in different order. During the whole task, all plasticity rules are frozen.

#### Training Phase

A layer *A* of linear readout neurons was trained in the following supervised way: We freeze the networks’ plasticity rules and generate an input timeseries 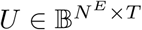 consisting of *T* = 10, 000 stacked input vectors ***u***(*t*) by repeatedly showing the sequences *s*_1_ and *s*_2_ in random succession. We record the stimulus-evoked activity timeseries 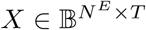 and the stimulus idendity 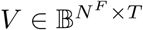. These timeseries consist of *T* = 10, 000 activity- and stimulus-identifier-vectors ***x***(*t*), ***v***(*t*), respectively. The timeseries were embedded with 0-vectors to achieve an offset of one timestep:

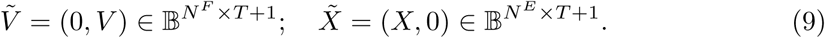

We want to predict ***v***(*t* + 1) from ***x***(*t*), so we have to find a matrix A such that

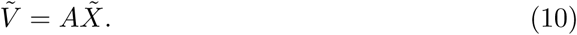

This equation is not strictly invertible for a non-quadratic matrix 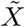, so we minimize the L2 norm:

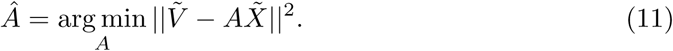

This minimization was computed using the python function numpy.linalg.lstsq and returned the readout layer *Â* as the result of the training phase.

#### Testing Phase

During testing phase, the performance of the readout layer *Â* was assessed on an independent input timeseries 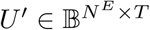 consisting of the same sequences *s*_1_,*s*_2_, but in different order. We now use the readout layer Â from the training phase to compute predictions 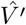 for the next letter of the independently generated timeseries *V′* from the testing phase:

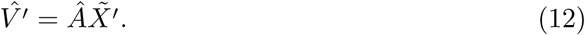

Afterwards, the performance *P* is computed as the number of successes in predicting the last letter of each sequence (either *c* or *f*). Thus, performance was computed as the difference of the third and the last component of the predicted identifier vector 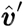 and real identifier vector 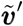, averaged across the whole duration of the testing phase *T* :

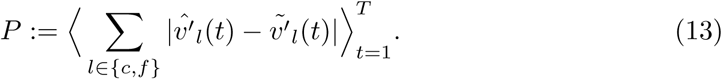

Note that this gives *P* ∈ [0, 1] and 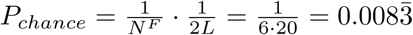. When scaling up the network size (Fig. 2-S1A, Fig. 3-S3A), we averaged over the prediction of all positions that were not random (*l* ∈ {*b, c, e, f*}) for increased stability of the performance measure.

### Frobenius Asymmetry Norm

During development, STDP increases the antisymmetry of the excitatory-excitatory connectivity matrix *W*^*EE*^ every time it acts. To measure the impact of STDP under the influence of SN, we first computed the Frobenius norm ∥ · ∥ _*ℱ*_ of the off-diagonal difference matrix *W*^*EE*^ − (*W*^*EE*^)^*†*^:

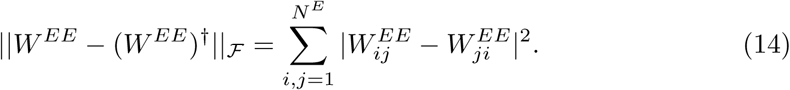

Here, (*W*^*EE*^)^*†*^ denotes the transpose. Then, we defined the antisymmetry 𝒜(*W*^*EE*^) over time as:

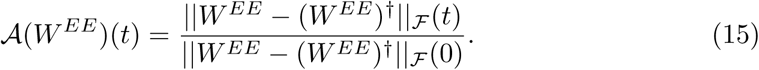

### Estimating mutual information

Mutual information was used to directly quantify how changes in the network architecture affect the capacity of neural activity to store past information. In general, the mutual information *I*[*Y, Z*] between two random variables *Y, Z* can be calculated from the marginal and joint probability distribution:

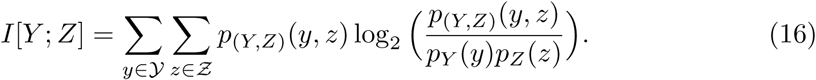

Specifically, we calculated the lagged mutual information *I*[*x*_*i*_(*t*); *v*_*j*_(*t* − *k*)] between the activity *x*_*i*_(*t*) of neuron *i* at time *t* and the presentation *v*_*j*_(*t* − *k*) of stimulus *j* at time *t* − *k*. This lagged mutual information provides an upper bound of the information about past stimuli *v*_*j*_(*t* − *k*) that a perfect decoder could read out when observing the activity *x*_*i*_(*t*) of a single neuron over one timestep. For the estimation of *I*[*x*_*i*_(*t*); *v*_*j*_(*t* − *k*)], stationarity and ergodicity were assumed, so that we can drop the time dependence and estimate 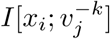 directly from a long simulation run of *T* time steps. The marginal (joint) probability distributions were approximated by counting how often the events occured during the long simulation run. To ensure that this estimation is accurate, a long simulation run of *T* = 100, 000 time steps in a frozen network driven by sequence input was used. Mutual information was then calculated using the `mutual_info_score` function in the `sklearn.metrics` python package.

### Calculating the Hamming distance

To assess the network dynamics in relation to criticality, we calculated the Hamming distance *D* as single-step approximation of the effective Lyapunov exponent [Lazar et al., 2009]. It allows the definition of three distinct dynamical regimes *D >* 1 marks the supercritical, unstable regime; *D <* 1 marks the subcritical, stable regime; and *D* = 1 marks the critical point. The Hamming distance *D* is calculated in a network under noise input, with frozen plasticity rules.

To examine network dynamics in isolation, each neuron’s activity in the absence of external input *ξ*_*i*_(*t* + 1) is calculated according to equation (1) with *u*_*i*_ = 0:

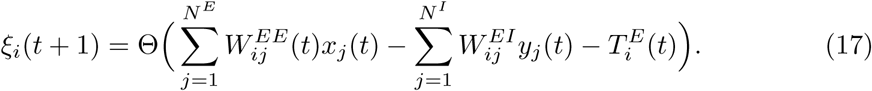

To test stability against internal perturbations, *N*^*E*^ hypothetical activities 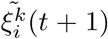 are calculated from *N*^*E*^ alternative pasts 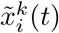. The alternative pasts are obtained by flipping the activity of unit *x*_*k*_(*t*), *k* ∈ {1, *…, N*^*E*^} from 0 to 1 or vice versa. The resulting hypothetical activities 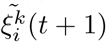 are then again calculated in the absence of external input:

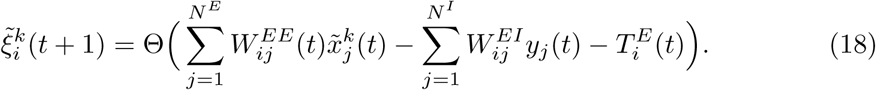

Next, we sum up the absolute distance between real *ξ*_*i*_(*t*) and hypothetical activities 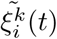 for all *N*^*E*^ neurons and average over the *N*^*E*^ alternative pasts:

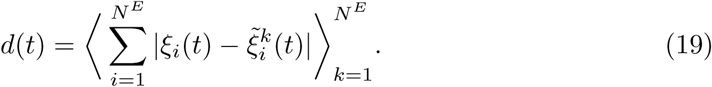

This average absolute distance *d*(*t*) is calculated for *T* = 100, 000 steps of network dynamics under noise input (plasticity rules frozen), and averaged over all timesteps, yielding the Hamming distance *D*:

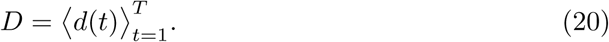

### Follower-sorting of the synaptic matrix

The synaptic matrix of the recurrent excitatory-excitatory connections *W*^*EE*^ was sorted along strongest synapses to reveal preferential paths. In detail, we constructed a linear transform from unordered “neuron id” space to “sorted id” space, which are identical vector spaces apart from the sorting of the dimensions. The first neuron in “neuron id” space was chosen as the first neuron (which is a random neuron because “neuron id” space is unordered) and the postsynaptic neuron of its strongest outgoing synapse was chosen as its “follower”, designated as the second neuron in “sorted id” space. This procedure was repeated starting from the former follower as the current neuron, choosing the postsynaptic neuron of its strongest outgoing synapse as the next follower, designating it as the third neuron in sorted id space etc. In the case that the postsynaptic neuron of the current neuron’s strongest outgoing synapse had already been designated above, the postsynaptic neuron of the second-strongest outgoing synapse was chosen as follower etc. This procedure reveals preferential paths of strong synapses on the secondary lower diagonal of *W*^*EE*^ in “sorted id” space.

### Calculation of the lag-one activation matrix

In order to show the presence of endogenous sequences in the spontaneous activity, the lag-one activation matrix *Q*^*EE*^ was calculated. In detail, the network activity under noise input was simulated for *T* = 10, 000 steps with frozen plasticity rules, and the spontaneous activity 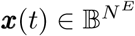 recorded. The lag-one activation matrix 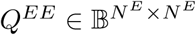 was now calculated as the tensor product of ***x***(*t*), ***x***(*t* − 1), corrected for the mean firing rate of the individual neurons and averaged over *T* :

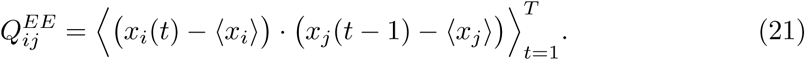

Here, 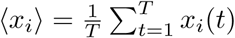 denotes the time average. Displaying *Q*^*EE*^ in the “sorted id” space obtained for *W*^*EE*^ reveals endogenous sequences on the lower secondary diagonal of *Q*^*EE*^.

### Implementation

The implementation of the model described above and the simulations presented in section “Results” where performed in the python programming language. The simulation and analysis codebase as well as all data necessary for reproduction of the figures presented here will be made available online (*https://github.com/Priesemann-Group*) upon publication.

## Acknowledgements

All authors received support from the Max-Planck-Society. Matthias Loidolt and Lucas Rudelt acknowledge funding by SMARTSTART, the joint training program in computational neuroscience by the VolkswagenStiftung and the Bernstein Network.

## Author contributions

Matthias Loidolt: Conceptualization, Methodology, Software, Formal analysis, Investigation, Data curation, Writing—original draft, Writing—review and editing, Visualization; Lucas Rudelt: Conceptualization, Methodology, Writing—review and editing, Supervision; Viola Priesemann: Conceptualization, Methodology, Formal analysis, Resources, Writing—review and editing, Supervision, Project administration, Funding acquisition

## Supplementary Figures for Main Text

**Figure 2-S1.**
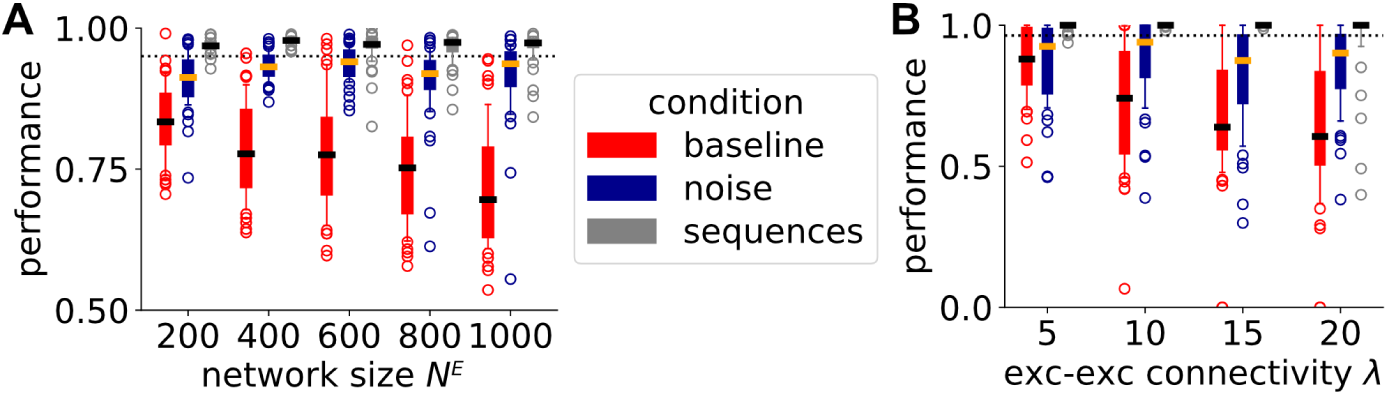
Improved sequence memory after development under noise in larger and denser networks. This figure extends Fig. 2 in the main text, comparing the same conditions in larger and denser networks. **A, B**. Development under noise (blue) increases sequence memory performance above baseline performance of the random initial network (red) for all network sizes (A) and all initial excitatory-excitatory connectivities (B) - but it never exceeds performance of networks developing under sequences (grey). Performance was measured after development for *t* = 20000 steps under either noise or sequence input. **A**. Influence of network size on sequence memory performance. Number of input units |*u*| and mean number of excitatory-excitatory connections *λ* were scaled with network size *N* ^*E*^ such that |*u*|*/N* ^*E*^ = 1*/*20 and *λ/N* ^*E*^ = 0.025, task difference was scaled up along network sizes: *L* = (10, 11, 12, 13, 14) for *N* ^*E*^ = (200, 400, 600, 800, 1000). **B**. Influence of excitatory-excitatory connectivity *λ* on sequence memory performance. Number of input units |*u*| = 10, network size *N* ^*E*^ = 200 and length of sequence *L* = 10 for all connectivities. Boxplots: Boxes extend from 25th to 75th quantile, whiskers from 10th to 90th quantile. Thick bar inside the box denotes the median, colored circles show outliers. Distributions were estimated from *N* = 50 independent realizations for each condition.

**Figure 3-S1.**
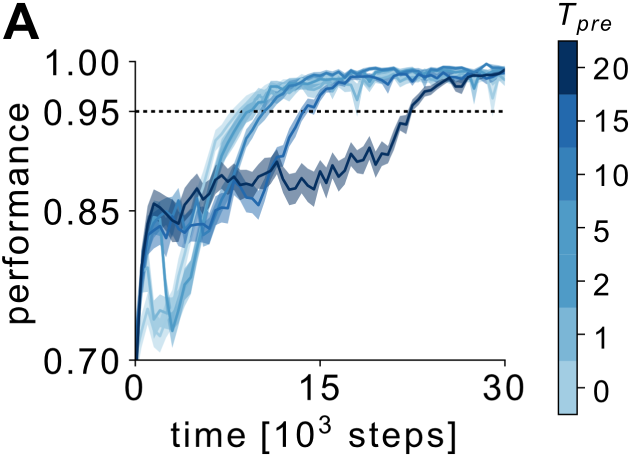
Learning from noise is slower than learning from sequences when including pre-shaping period. This figure shows the same data as Fig. 3 in the main text, but shows them on a different abscissa using absolute time. **A**. Two-phased development process with different inputs: First, development under noise for different pre-shaping periods *T*_*pre*_. Second, development continues after onset of sequence input at *t* = *T*_*pre*_. During development under noise, networks show a faster increase up to *P* = 0.85 but are slower to reach the *P* = 0.95 threshold than networks starting with sequence input, regardless of *T*_*pre*_. Lines: mean over *N* = 50 independent realizations; shaded areas: 33 − 66 % confidence interval thereof.

**Figure 3-S2.**
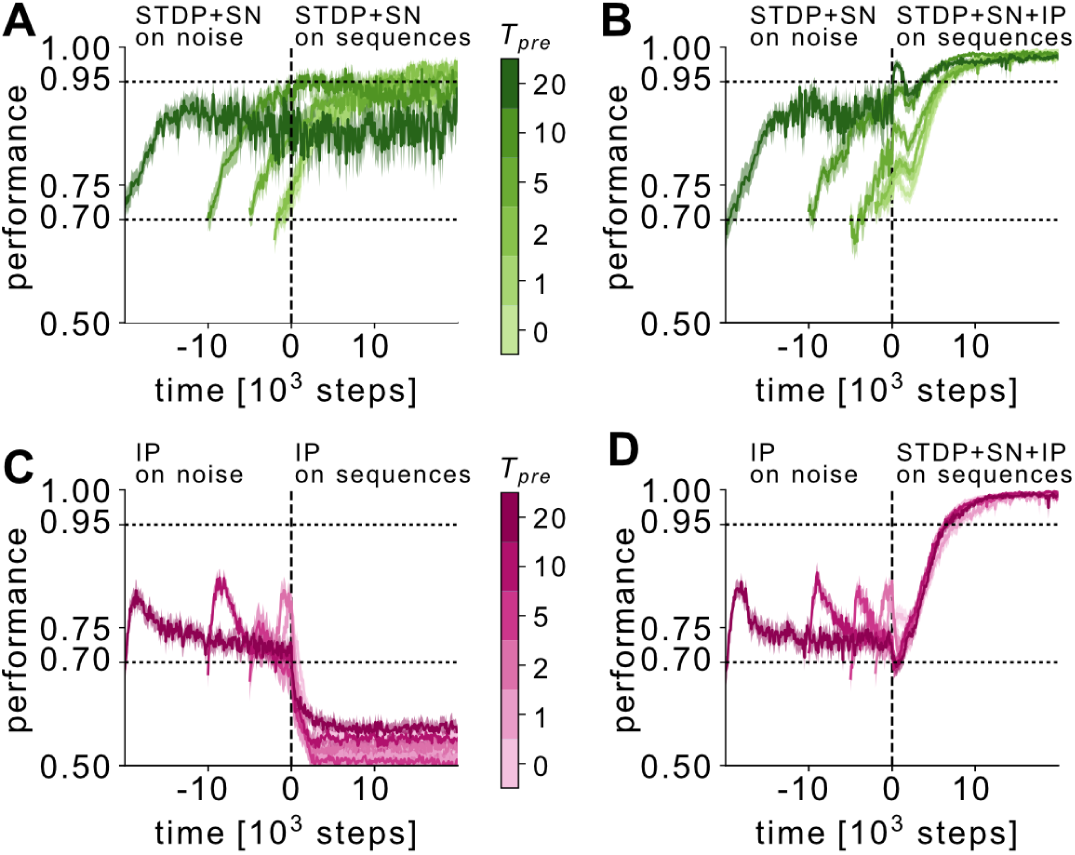
Pre-shaping under noise requires both spike-timing dependent and intrinsic plasticity. This figure corresponds to Fig. 3 in the main text, but does not use all different plasticity rules all the time. To isolate the pre-shaping effects of the individual plasticity rules, we modified the two-phased development to start with either spike-timing dependent plasticity and synaptic normalisation (STDP+SN) only (A,B) or intrinsic plasticity (IP) only (C,D). We then continued development under the same plasticity rules (A and C, respectively) or under all plasticity rules (B and D, respectively) to separate the effects of sequence input on each plasticity rule from the pre-shaping effect of each plasticity rule. **A**,**B**. Two-phased development process with STDP+SN(+IP) plasticity rules: first, development under noise and STPD+SN for different pre-shaping periods *T*_*pre*_. Second, development continues after onset of sequence input, under (A) STDP+SN and (B) STDP+SN+IP, respectively. **A**. Pre-shaped networks (*T*_*pre*_ *>* 0) maintain higher sequence memory performance than those starting from initial random conditions (*T*_*pre*_ = 0), but do not reach maximum performance. **B**. Pre-shaped networks (*T*_*pre*_ *>* 0) maintain higher sequence memory performance than those starting from initial random conditions, but show a transient dip in performance before reaching the maximum. **C**,**D**. Two-phased development process with IP(+STDP+SN) plasticity rules: First, development under noise and IP for different pre-shaping periods *T*_*pre*_. Second, development continues after onset of sequence input, under (C) IP and (D) IP+STDP+SN. **C**. Pre-shaped networks (*T*_*pre*_ *>* 0) show transient peak in performance during noise phase before relaxing back to baseline. Upon sequence input, performance drops below baseline regardless of the length of the pre-shaping period *T*_*pre*_. **D**. Pre-shaped networks (*T*_*pre*_ *>* 0) show transient peak in performance during noise phase before relaxing back to baseline. Upon sequence input, performance increases to the maximum, in a parallel fashion regardless of the length of the pre-shaping period *T*_*pre*_. Lines: mean over *N* = 50 independent realizations; shaded areas: 33 − 66 % confidence interval thereof.

**Figure 3-S3.**
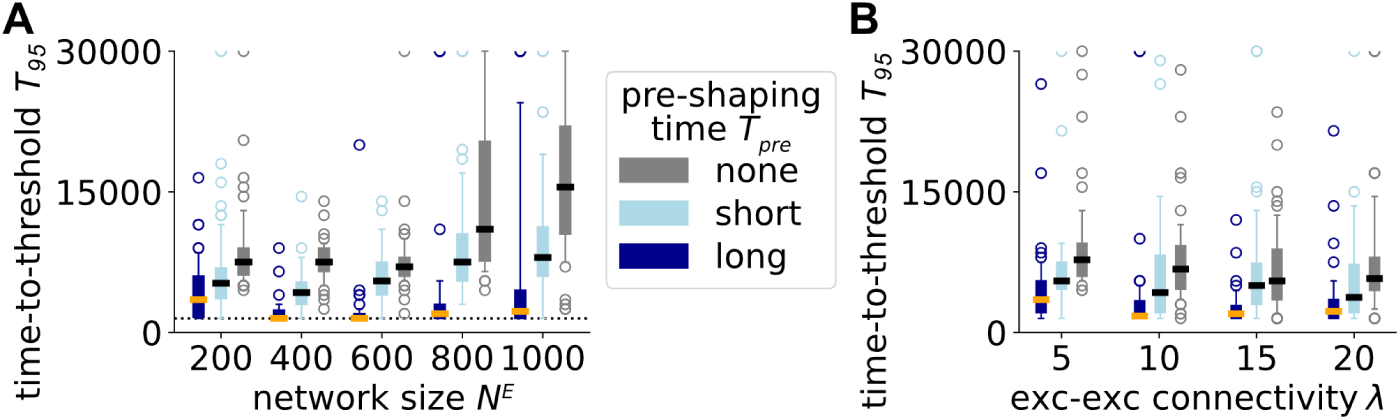
Faster sequence learning after noise pre-shaping in larger and denser networks. This figure extends Fig. 3 in the main text, comparing the same conditions in larger and denser networks. **A**,**B**. Networks pre-shaped by development under noise (blue) learn specific sequences faster than the random initial network (grey), regardless of network size or initial connectivity. The panel depicts the time of development after sequence onset that is required to reach a 95 % performance threshold. **A**. Influence of network size on pre-shaping efficiency. Number of input units |*u*| and mean number of excitatory-excitatory connections *λ* were scaled with network size *N* ^*E*^ such that |*u*|*/N* ^*E*^ = 1*/*20 and *λ/N* ^*E*^ = 0.025, task difference was scaled up along network sizes: *L* = (10, 11, 12, 13, 14) for *N* ^*E*^ = (200, 400, 600, 800, 1000). Performance was measured after development under sequences for *t* = 15000 (light blue) and *t* = 30000 (dark blue). **B**. Influence of excitatory-excitatory connectivity *λ* on pre-shaping efficiency. Number of input units |*u*| = 10, network size *N* ^*E*^ = 200 and length of sequence *L* = 10 for all connection densities *λ*. Performance was measured after development under sequences for *t* = 5000 (light blue) and *t* = 15000 (dark blue). Boxplots: Boxes extend from 25th to 75th quantile, whiskers from 10th to 90th quantile. Thick bar inside the box denotes the median, colored circles show outliers. Distributions were estimated from *N* = 50 independent realizations.

**Figure 3-S4.**
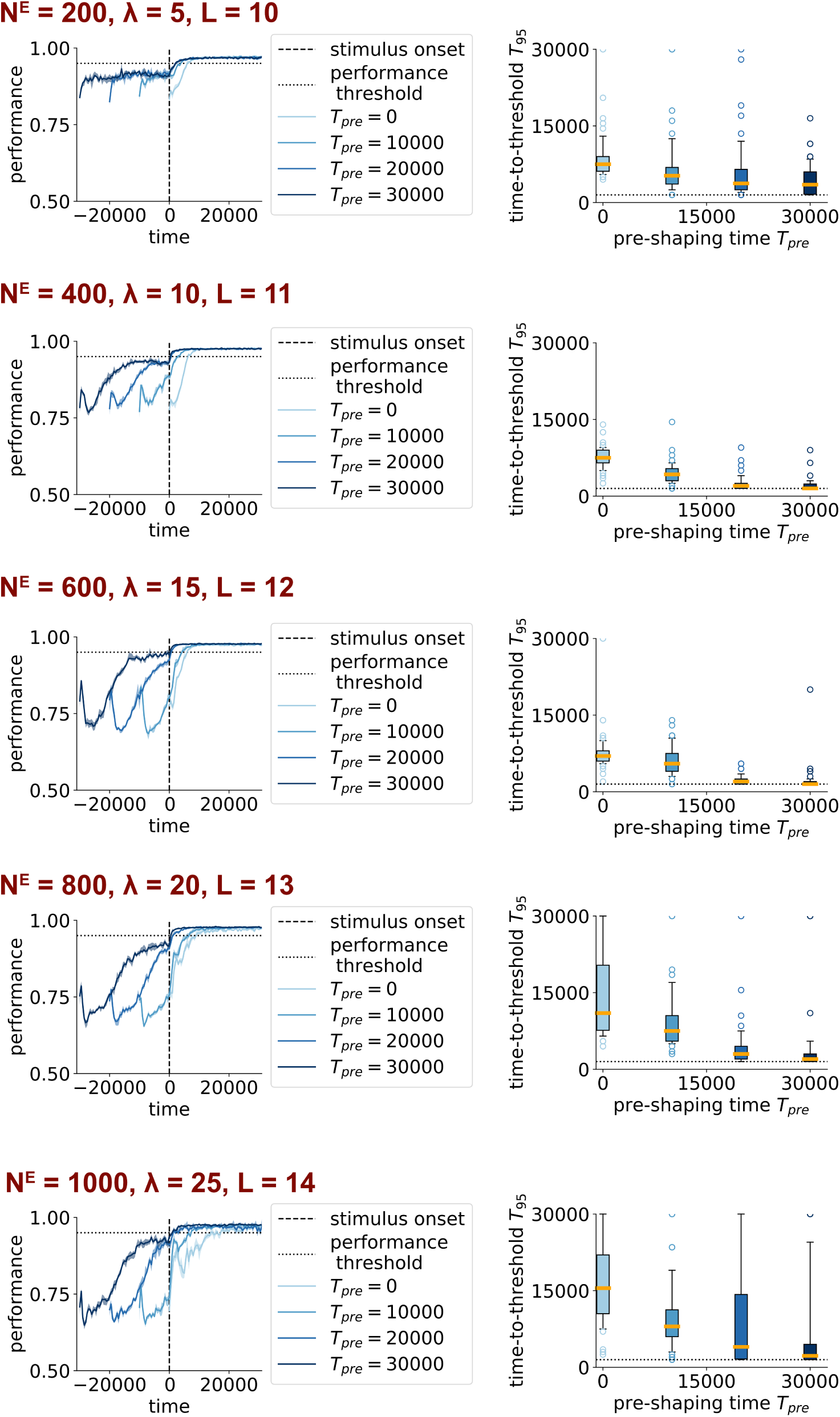
Improved sequence memory and faster sequence learning after noise pre-shaping in larger networks. This figure corresponds to Fig. 3 in the main text, showing the same scenario for networks of increasing size. *N* ^*E*^ denotes the number of excitatory neurons, *λ* the mean number of exc-exc connections, and *L* the length of the sequence that has to be memorized. Note that *L* has to be increased with *N* ^*E*^, otherwise the task would become too simple. Lines: mean over *N* = 50 independent realizations; shaded areas: 33 − 66 % confidence interval thereof.

**Figure 3-S5.**
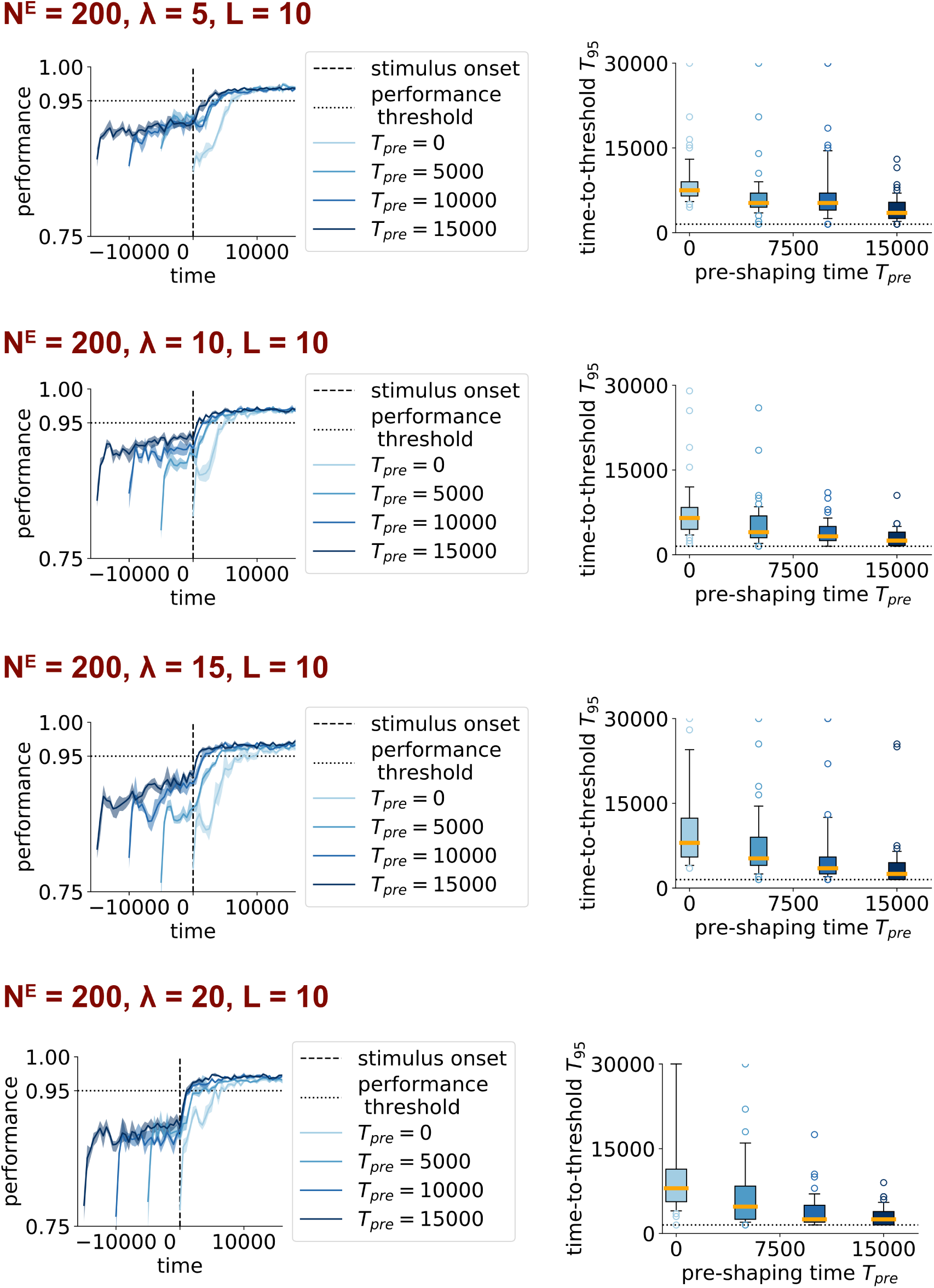
Improved sequence memory and faster sequence learning after noise pre-shaping in denser networks. This figure corresponds to Fig. 3 in the main text, showing the same scenario for networks of increasing density. *N* ^*E*^ denotes the number of excitatory neurons, *λ* the mean number of excitatory-excitatory connections, and *L* the length of the sequence that has to be memorized. Note that *L* does not change with *λ*, i.e. denser networks can perform similarly hard tasks. Lines: mean over *N* = 50 independent realizations; shaded areas: 33 − 66 % confidence interval thereof.

**Figure 5-S1.**
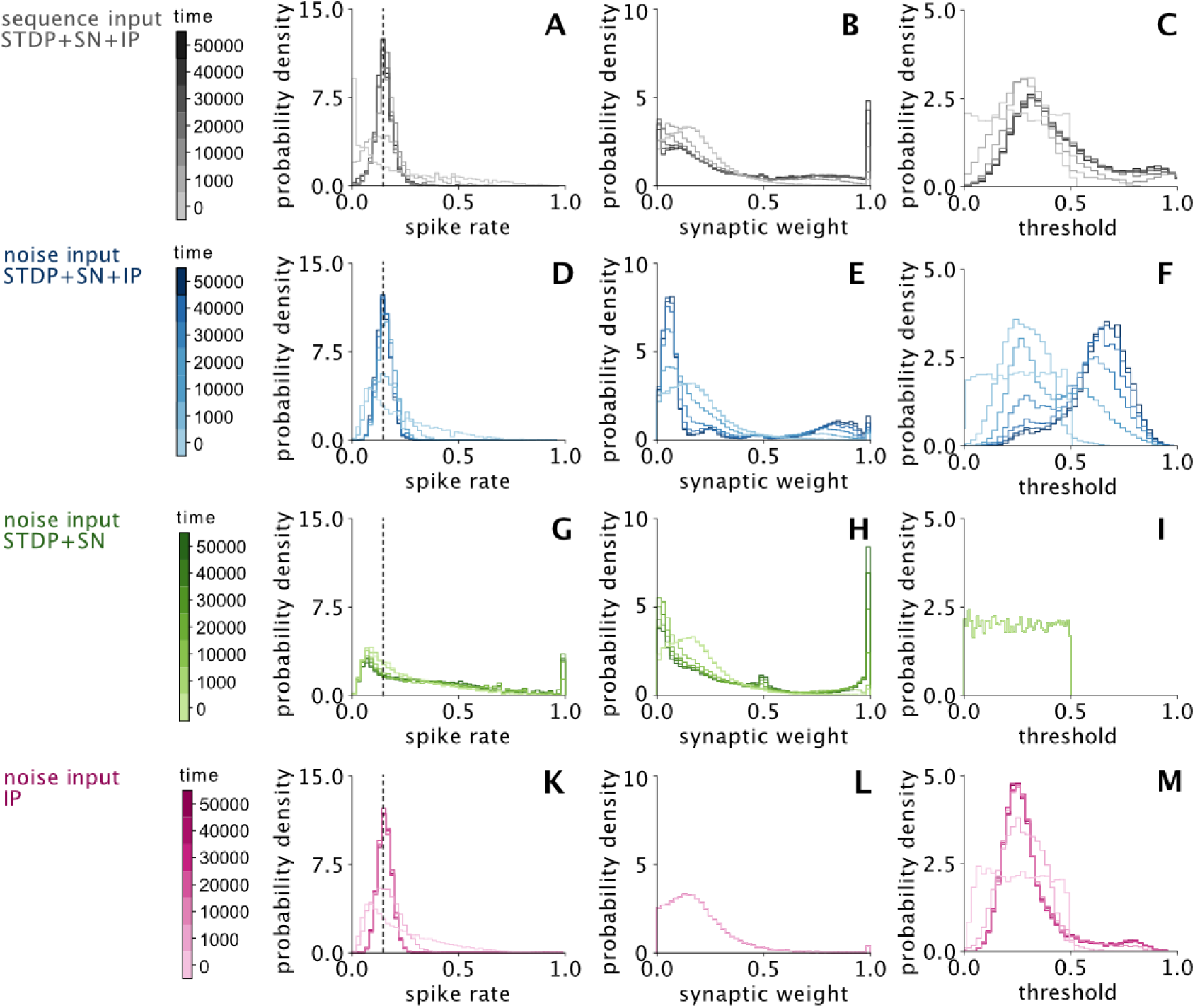
All plasticity rules are necessary to ensure physiological activity and long-tailed synaptic efficacy distributions. This figure extends Fig. 5 in the main text, showing the network structure throughout development. **A**.,**D**.,**G**.,**K**. Spike rate distributions. IP quickly ensures a close-to normal firing rate distribution centered on the target rate *r*^*^ marked by the dashed line (A, D, K). The absence of IP leads to many neurons being less active than the target rate, while some have unphysiologically high spike rates (G). **B**.,**E**.,**H**.,**L**. Synaptic weight distributions. STDP+SN selectively strengthens synapses, either because of structured sequence input or because of high initial weight, resulting in a bimodal weight distribution (B, E, H). In the absence of IP, the unregulated winner-take-all dynamics caused by STDP+SN grow a large proportion of synapses to have a weight of unity (H). **C**.,**F**.,**I**.,**M**. Threshold distributions. IP quickly adapts the random initial uniform distribution, silencing neurons receiving excess sequence input (long tail in C) and allowing for participation of all other neurons (C, F, M). Under noise input, strong input is more equally distributed, so IP slowly increases the mean threshold to balance changes made by STDP+SN (F). In the absence of STDP+SN, IP slowly changes thresholds to match the initial weight distribution (shown in L), causing the tailed distribution seen in (M). All lines show histograms obtained by pooling over *N* = 50 random initial realizations.

## Supplementary Results

### Intrinsic plasticity causes fast performance increase under noise input but will threshold preferential paths in isolation

IP alone suffices to reproduce the rapid rise but can not sustain high performance for more than *τ*_*IP*_ steps (Fig. SR-1A): Surprisingly, the performance reached during this rapid, transient rise in the noise phase will break down immediately upon onset of sequence input if the effects of STDP+SN have not yet materialized (*t < τ*_*ST DP*_). This immediate breakdown in performance must also be caused by IP alone because STDP+SN is not fast enough to act on this timescale.

The rapid, but unsustained increase of sequence performance under noise input is due to the initial threshold distribution not being optimal. Under noise input (uniformly distributed external drive), the spike rate of each neuron reflects the mismatch between its threshold and the recurrent drive it receives. This mismatch is quickly corrected by the IP rule: it steers each neurons spike rate towards target rate, enabling close-to-optimal performance for the initial random matrix (Fig. SR-1A,B). Yet, this performance slowly decays again because the threshold distribution continues to develop even after the network target rate is reached (Fig. SR-1C).

Both the slow performance decay after *t > τ*_*IP*_ in the “noise IP only” case and the breakdown seen upon switching to sequence input after short development under noise (*T*_*pre*_ *< τ*_*ST DP*_) are due to IP thresholding preferential paths of activity. Neurons with a dominant incoming synapse are slowly silenced by IP under noise input after the network activity approaches target rate (Fig. SR-1C,D), because the combined drive from the strong synapse and the external input slightly exceeds the target rate. In contrast, IP begins to silence neurons preferentially activated by sequence input right from sequence onset, rapidly silencing the preferential paths leading away from the input neurons (Fig. SR-1D). Shuffling the thresholds between neurons breaks these spatial dependencies and rescues performance back to the level observed when target rate was first reached (Fig. SR-1A). Thus, both the slow decay of performance under noise input as well as the performance dip observed at sequence onset after short pre-shaping times are due to IP thresholding weak preferential paths crucial for sequence memory.

**Figure SR-1.**
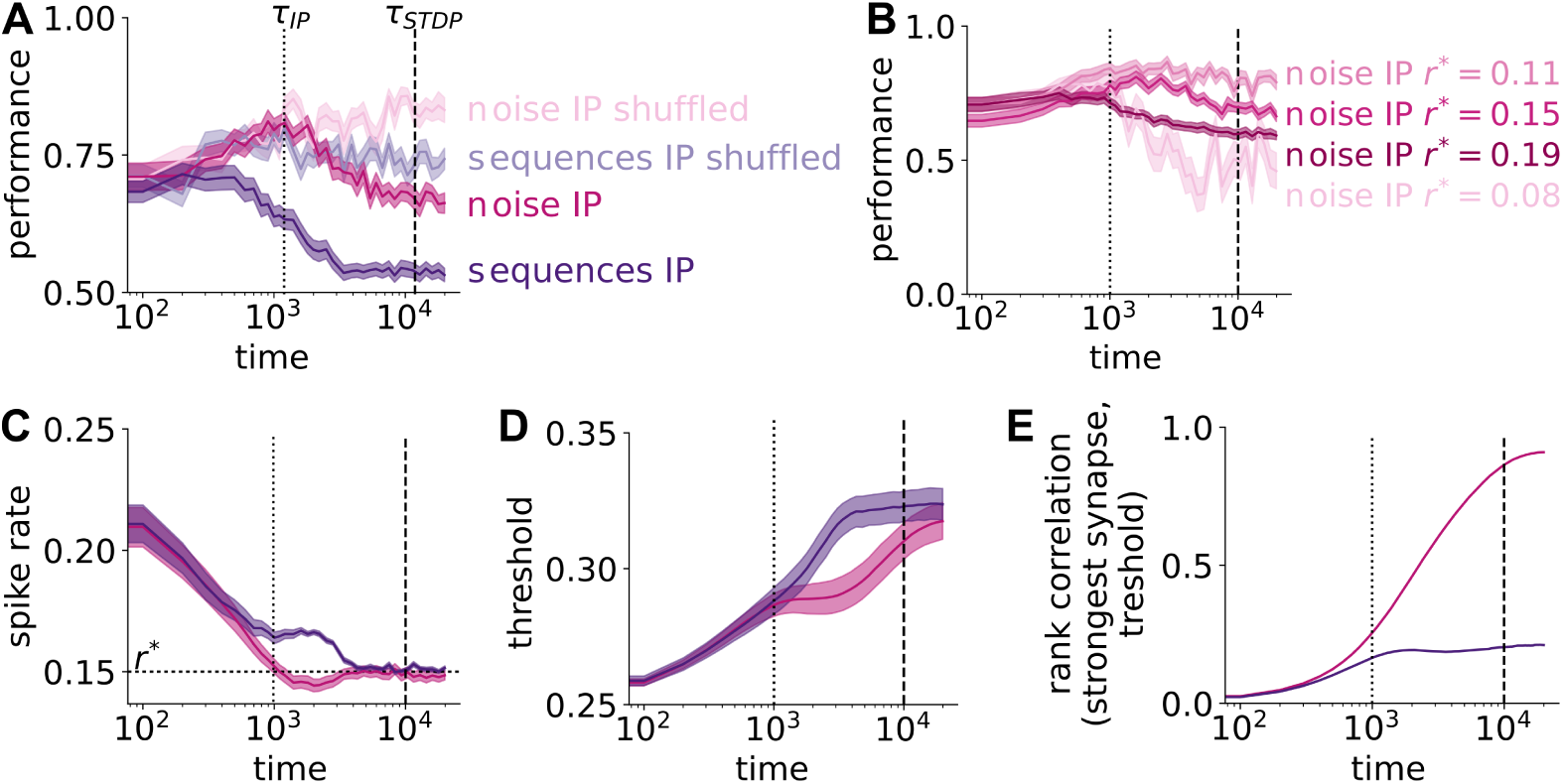
Intrinsic plasticity causes fast performance increase under noise input but will threshold preferential paths in isolation. **A**. Noise input and IP cause fast, but transient, increase in sequence memory performance. Under sequence input, IP causes an immediate breakdown in performance. Shuffling the thresholds adapted by IP, thus discarding spatial correlations between input structure and activity, restores performance. **B**. Performance peak caused by noise input and IP depends on target rate *r*^*^. Fine-tuning the target rate can increase the peak performance observed around *τ*_*IP*_, but the performance will eventually fade for all target rates. **C**. Noise input and IP (pink) cause fast decrease of network spike rate because all neurons receive a similar external drive. Networks developing under sequence input (violet) take longer than *t > τ*_*IP*_ to reach the target rate. This is due to some neurons (those stimulated by the letters *b*; *d*) receive external drive exceeding the target rate, necessitating strong threshold growth to reach target rate. **D**. Under noise input, the mean threshold develops in two phases: first, it generally increases such that the network reaches target rate. As most neurons have reached target rate (at *t* ≈ *τ*_*IP*_ = 10^3^), those with a dominant incoming synapse increase their threshold further (until *t* ≈ 2 *·* 10^4^). Under sequence input, neurons that are preferentially recruited by the externally drives will be silenced within *t* ≈ 4 *·* 10^3^). **E**. Correlating the weight of the strongest incoming synapse and the threshold evidences that under noise input, the threshold is increased to match the strongest synapses. Under sequence input, those neurons are silenced which are preferentially activated by the external stimulation - hence correlation between strongest incoming synapse and threshold remains low. Target rate *r*^*^ = 0.15 in all panels except (B). Lines: mean over *N* = 50 independent realizations; shaded areas: 33 − 66 % confidence interval thereof.

